# A human intracranial map of consciousness returning from anesthesia

**DOI:** 10.64898/2026.06.17.732807

**Authors:** Xinyu Chen, Haoru Zhang, Xinran Deng, Yong Ren, Yiting Liu, Jialei Wang, Shiyi Xu, Yuchao Ji, Zihan Yang, Wenwen Jia, Xialei Huang, Yuanqing Wang, Lanhui Huang, Sixian Li, Yue Yuan, Aoqing Luo, Junfan Chen, Chen Yao, Yuchen Xiao

## Abstract

How consciousness arises from the brain remains a central yet unsolved question. Although substantial effort has been devoted to characterizing the neural correlates of conscious perception during wakefulness, how consciousness re-emerges from unconscious states remains poorly understood. Here, we took advantage of the rare opportunity to record human intracranial local field potentials from propofol-induced general anesthesia through the transition to behavioral engagement and subsequent full wakefulness, capturing both the stable states of unconsciousness and consciousness and the reconstituting neural dynamics that bridge them. Anesthesia cannot be defined by any single electrophysiological signature; rather, it is an organized low-frequency regime characterized by a constellation of coordinated neural phenomena, including aperiodic slow waves, alpha/beta periodicity, global alpha synchronization, and slow-wave-alpha/beta phase-amplitude coupling. Following anesthetic cessation, this regime progressively dissolved as neural excitability and dynamical complexity climbed. The appearance of conscious behavior coincided with a fast and dramatic transformation in high-gamma activity, from stochastic and unpredictable bursts to structured responses that were distributed, task-selective, and event-locked. These findings suggest that the recovery of consciousness is a multiscale reorganization in which distinct neural underpinnings rise and fall. Together, this work charts an electrophysiological map of how conscious cognition is extinguished, reconfigured, and restored in the human brain.

## Introduction

The neural fabric of consciousness is arguably one of the most intriguing scientific quests today. Once marginalized as elusive, subjective, and ill-defined for rigorous study, consciousness has now earned its position in serious scientific investigation. Conscious access^1–3^ can be reliably manipulated through deft experimental paradigms including binocular rivalry^4^ and continuous flash suppression^5^, or through pharmacological agents such as anesthetics^6,7^ and psychedelics^8,9^. Advanced neurobiological^6,10–12^, neural recording^11,13,14^, neuroimaging^15,16^, and neuromodulatory^17,18^ tools have allowed for the identification of neural correlates of consciousness (NCCs). Theories of consciousness^1,3,19–24^, spanning physical, biological, and computational perspectives—often with substantial conceptual overlap—are already in place to be fortified or falsified. Together, these developments have transformed consciousness from a philosophical abstraction into a concrete scientific object.

A central question in this nascent domain is how consciousness arises. In addressing this challenge, general anesthesia presents distinctive advantages. First, it is unclear to what degree the prolonged unconscious state induced by anesthesia mechanistically resembles or differs from the lack of conscious access to specific contents during wakefulness^25^, as in continuous flash suppression. Overwhelmingly more experiences are abolished by general anesthesia^25,26^. Certain anesthetics including propofol and xenon are thought to induce states with little or no conscious experience, while others like ketamine may generate dream-like experiences where there is “disconnected consciousness”^26,27^. We believe that identifying the neural correlates of unconsciousness (NCUCs) is as equally critical as searching for the NCCs en route to demystifying consciousness. Second, the emergence from anesthesia reflects a continuous reconstruction of consciousness from its loss, a process that is unattainable through paradigms in which conscious access swiftly switches between all and none. Therefore, examining the neural processes from anesthesia to wakefulness yields valuable insights into how consciousness is suppressed and generated within the brain, and directly informs the understanding of the neurobiology of consciousness and anesthesia itself^6,22,26^.

The pharmacological and molecular actions of commonly used anesthetics have been extensively studied, largely through animal and in vitro investigations^28–30^. There have been attempts to examine consciousness in animals using anesthesia^31–34^; however, the absence of subjective reports makes it challenging to infer the precise level or content of consciousness from behavior alone^35^. Moreover, the nature of consciousness across species might be fundamentally discrepant^36^. Most human studies that borrow anesthesia for investigating consciousness have relied on non-invasive measures like scalp-EEG^37–39^ and fMRI^40,41^. These methods, despite their advantages of broad cortical coverage and applicability to healthy subjects, yield readouts with relatively limited spatiotemporal resolution and do not adequately sample deep brain structures^42,43^. Studies combining human intracranial recordings with anesthesia to directly investigate consciousness remain scarce. In addition, most previous intracranial studies have focused on circumscribed anatomical regions or selected anesthetic stages, often examining only one or two isolated neural phenomena under task-free or simplified-task settings^44–51^. Thus, what remains to be established is a regionally comprehensive and temporally continuous account of how the brain progresses from anesthetic unconsciousness to the restoration of conscious cognition, while characterizing the reconstitution of distinct mental faculties.

In this study, we took advantage of the rare opportunity to invasively record human neural electrophysiological signals during propofol-induced anesthesia and emergence. Subjects were patients with pharmacologically intractable epilepsy, who received stereo-EEG electrode implantation surgery for localizing epileptogenic foci. A multimodal task assessing basic auditory, motor, communicational, and visual functions were administered continuously and repeatedly throughout the session, spanning states from general anesthesia to full wakefulness. By probing neural dynamics across this trajectory, we provide a systems-level framework for understanding how consciousness is extinguished, reconstituted, and restored in the human brain.

## Results

Intracranial local field potentials (LFPs) were recorded in the operating room (OR) where patients with pharmacologically intractable epilepsy received stereo-EEG electrode implantation surgery for seizure localization. We obtained LFPs from a total of 1190 channels (**Figure 1d, Tables S1–2**) in 10 subjects (**Table S3**). Recordings began during propofol-maintained general anesthesia and continued until patients achieved sustained perfect performance across the multimodal task (**Figures 1a–b**). One or two days after the implantation surgery, patients performed the same task again while wide awake in the epilepsy monitoring unit (EMU). The results are presented chronologically, following the sequence of events in the OR and then the EMU, with behaviors and their neural correlates described last.

**Figure 1.**
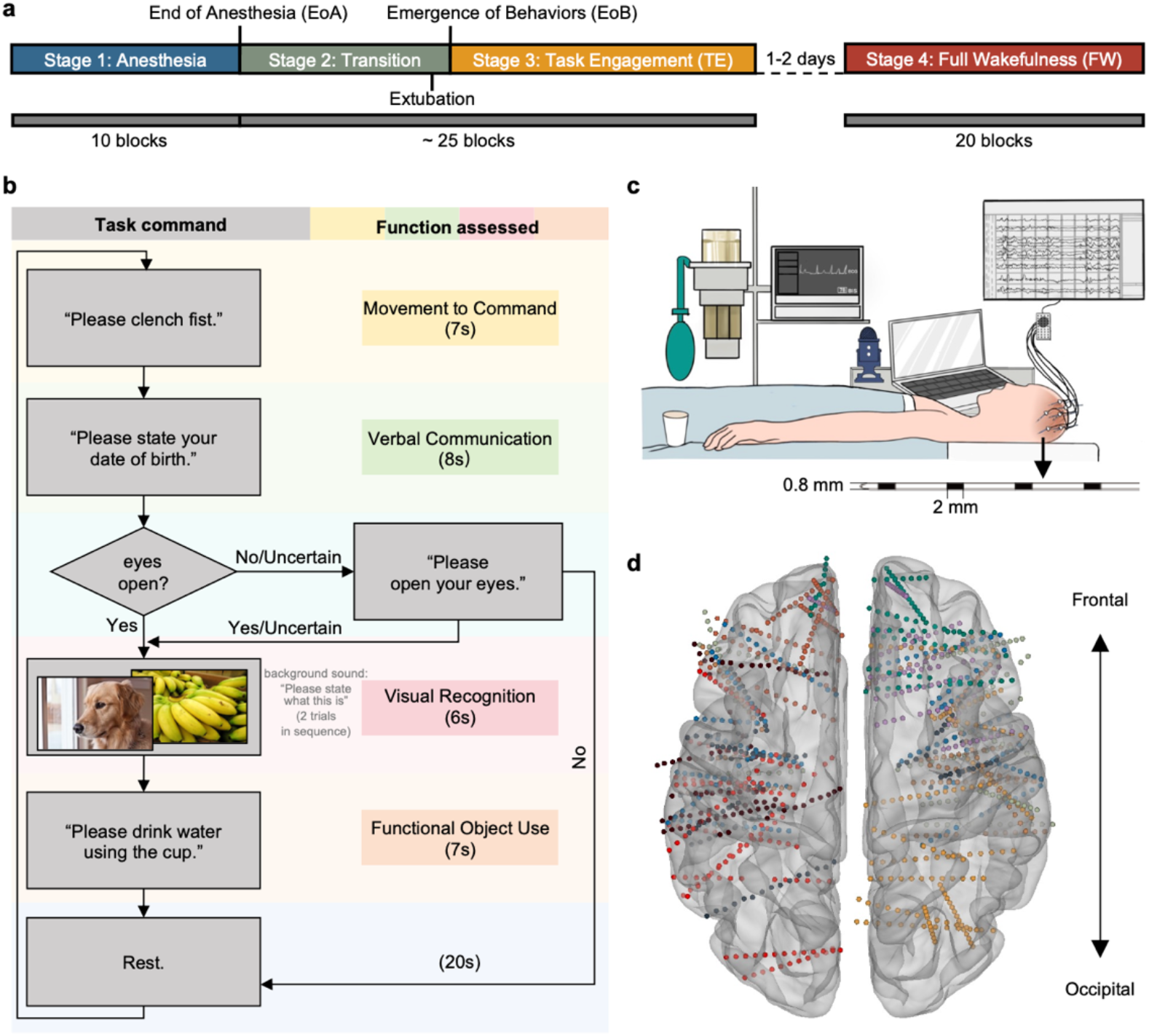
Methods synopsis. **a**. General procedures. The recording contained four stages: Anesthesia (Stage 1), Transition (Stage 2), Task Engagement (TE, Stage 3), and Full Wakefulness (FW, Stage 4). Stages 1–3 took place in the OR, whereas Stage 4 in the EMU. Stages 1–3 were demarcated by two key events: EoA and EoB. Extubation typically occurred a short time before EoB. After one or two days, the same task was administered again (Stage 4). **b**. Each block comprised two trials before eye opening and four trials after eye opening, with a 20 s inter-block rest period. **c**. Device and equipment setup. Each stereo-EEG channel has a length of 2 mm and a diameter of 0.8 mm. **d**. Anatomical locations of all channels (bottom view). Different colors denote channels from different subjects.

### Prominent electrophysiological changes after the End of Anesthesia and before the Emergence of Behaviors

Time-frequency representation revealed LFP power changes around key events: EoA, Extubation, and EoB (**Figure 2a**). We performed an RBF (radial basis function)-based change point (CP) detection (**Methods**) to pinpoint the timings of abrupt changes in the electrophysiological profile. RBF-kernel CP detection can identify nonlinear and higher-order changes in the joint distribution of multivariate signals beyond simple shifts in mean or variance, rendering it particularly effective for neural electrical signals with intricate correlational and high-dimensional dynamics. **Figure 2b** depicts the locations of CPs identified from all channels in one subject, with density indicating the degree of CP concentration. The two clusters were sandwiched between EoA and EoB. Most channels exhibited two CPs from anesthesia to emergence (**Figures 2c** and **2f**). **Figures 2d–e** illustrate the distribution of CPs across all subjects, again revealing two clusters during Transition. These results suggest that neural electrophysiological activities underwent significant changes during the recovery from propofol-induced anesthesia, and remained changing until conscious behaviors surfaced.

**Figure 2.**
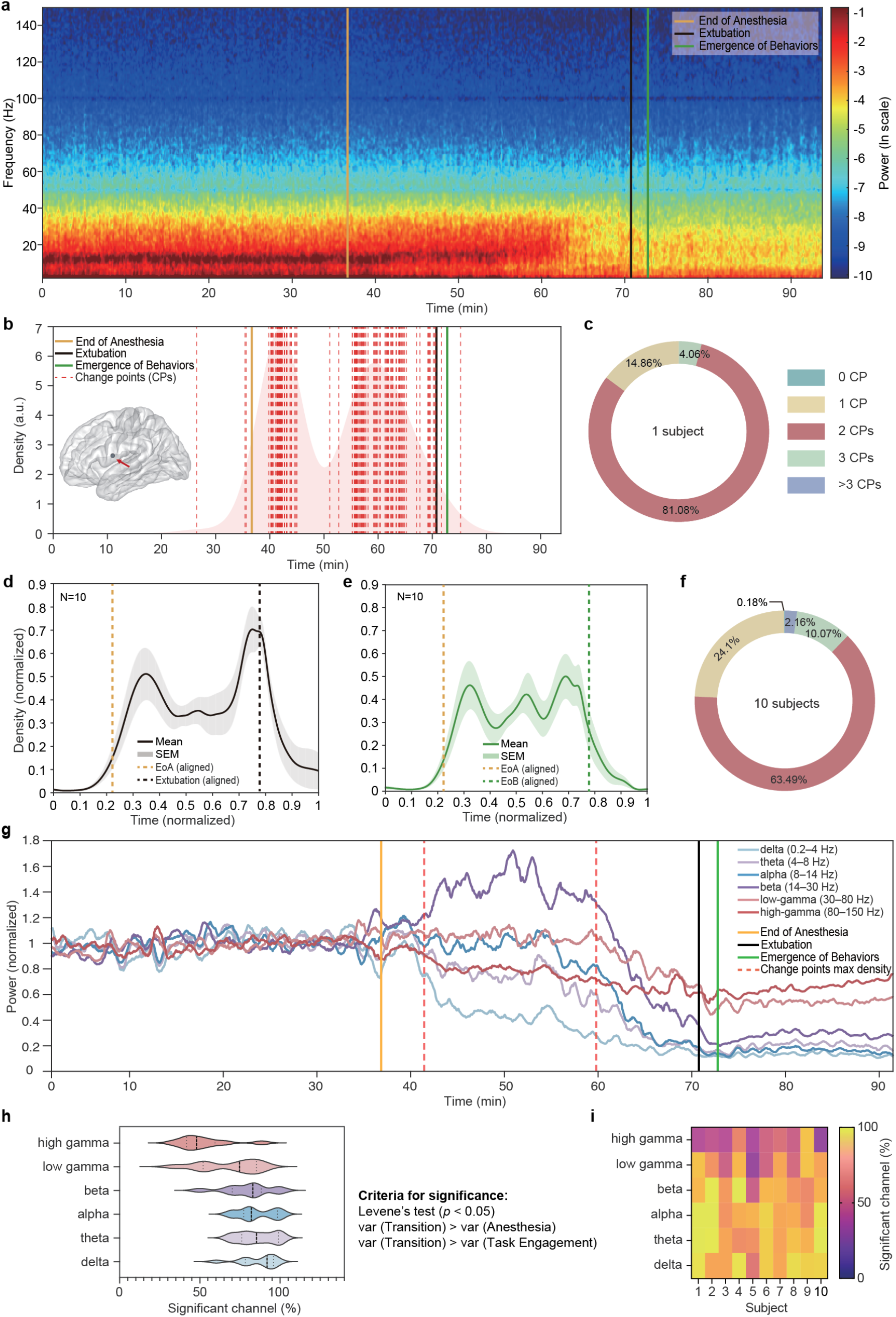
Substantial electrophysiological changes during Transition. **a**. Time-frequency decomposition of the LFP signal from one channel (precentral) in one subject. **b**. Distribution of CPs from all channels. Density indicates the degree of clustering. Inset shows the location of the channel in **a. c**. Grouping of channels by the number of CPs identified in each, showing results from the same subject in **b. d–e**. Averaged distribution of CPs across all subjects, temporally aligned to Extubation (**d**) or EoB (**e**). For each individual, the distribution was normalized such that *den*_*si*_*ty*_*min*_ = 0 and *den*_*si*_*ty*_*max*_ = 1. **f**. Grouping of channels from all subjects by the number of CPs identified in each channel. **g**. Frequency-specific power changes in one subject. Stages 1 and 3 were more stable than Stage 2. Each line represents the median power of a frequency band across all channels. Power values during Anesthesia were calibrated to 1 for better visualization of the change in variance over time. **h**. The proportion of channels that demonstrated higher variance in Transition, separated by frequency bands. **i**. Same data as in **h**, further separated by subjects.

Next, we performed time-frequency decomposition to zoom into the neural activities in each frequency band. **Figure 2g** shows an example of the temporal evolution of power across frequency bands in one subject. During Anesthesia and TE, power across all frequency bands was relatively stable, in sheer contrast to the intense fluctuations during Transition. The dashed lines correspond to the two CP density peaks shown in **Figure 2b**. The first peak marked a sharp increase in power variability, whereas the second forecasted its subsequent decline. Levene’s test identified channels whose power variance during Transition exceeded that observed in both Anesthesia and TE. This pattern was particularly prominent in lower-frequency bands (delta: 86.3%, theta: 85.6%, alpha: 85.5%, beta: 79.7%; mean: 84.3%; **Figure 2h**). Although less pronounced at higher frequencies, elevated variance during Transition was still observed in the majority of channels (low-gamma: 68.2%, high-gamma: 51.9%; mean: 60.1%). **Figure 2i** displays the results in each individual. Together, the clustered change points and elevated power variability reveal a stable–unstable–stable neural trajectory from anesthesia to emergence, suggesting that the most substantial electrophysiological reconfiguration occurs during the Transition period before but not after conscious behavior manifests.

### Rising excitation and complexity during consciousness return, with lower levels in the temporal lobe

Next, we sought to identify what underlay the remarkable electrophysiological changes during Transition. We first considered the excitatory-to-inhibitory (E/I) balance in the brain, given that propofol promotes the function of the inhibitory neurotransmitter GABA. We predicted an increase in E/I balance throughout the procedure, and that the beginning and the end of such this increase would align with the distribution of CPs. The E/I balance can be computationally estimated using electrophysiological signals^52^. Background electrical activity in the brain is regarded as aperiodic and can be described with 1/*f*^*x*^ (*f*: frequency; *x*: exponent) dynamics in the power spectrum^53^. Previous studies^53^ have demonstrated that 1/*x* can be a spectral proxy measure for E/I balance (**Methods**). **Figure 3a** illustrates an example of how 1/*x* shifted across stages in one subject. 1/*x* sharply increased during Transition and stabilized during TE. **Figures S1a–i** displays the results from all other individuals. Population results (**Figure 3b**) showed the same trend, where 1/*x* steadily climbed during Transition and plateaued in TE. These results are in conformity with the locations of CPs that were clustered within Transition (**Figure 2e**). The 1/*x* values were significantly different across the three stages both at the group level (**Figure 3c**) and at the single-subject level (**Figures S2a–j**). In all brain regions, 1/*x* was significantly higher during Task Engagement than during Anesthesia (**Figure 3g**).

**Figure 3.**
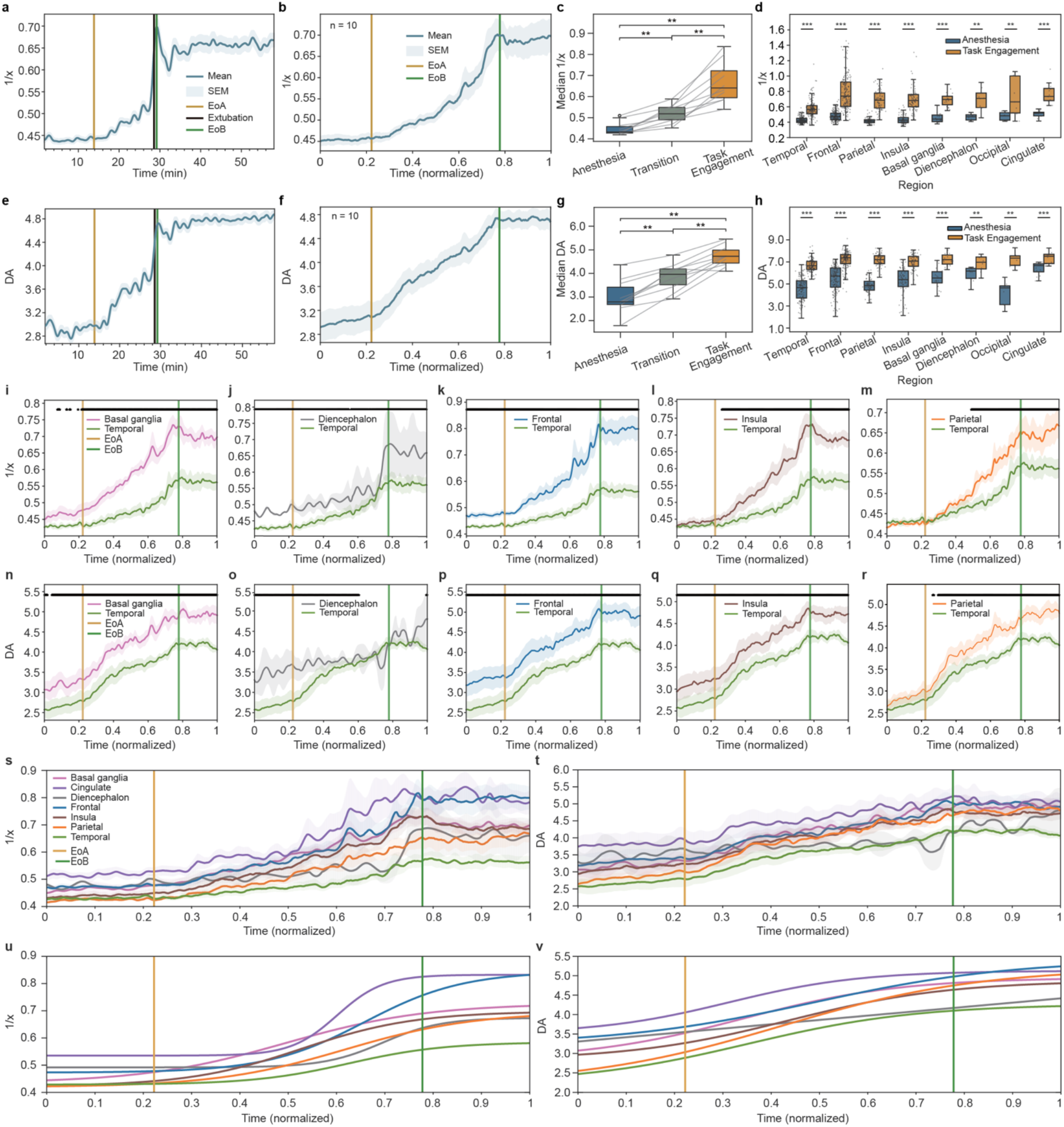
Increase of E/I balance and dimension of activation during Transition and their comparison among stages and regions. **a**. An example of how the E/I balance, estimated with 1/*x*, changed over time in a single subject. The main line indicates the averaged 1/*x* from all channels in this subject. **b**. Temporally aligned mean 1/*x* across all subjects. **c**. Comparison of 1/*x* across stages. Each line represents data from a single subject, connecting the median 1/*x* values across all channels within that subject. Pairwise comparisons were conducted using the signed-rank test. **d**. 1/*x* values across different regions during Anesthesia and TE. Signed-rank test was implemented for pairwise comparison in each region. **e–h**. Results of *DA* in the same format as in **a–d. i–m**. Comparisons of temporally aligned 1/*x* between the temporal lobe and other regions. Black horizontal lines indicate the time points that showed significant differences between regions (linear mixed model, FDR-corrected *p* < 0.05). **n–r**. Results of *DA* in the same format as in panels **i–m. s–t**. 1/*x* (**s**) and *DA* (**t**) trends in all regions. **u–v**. Sigmoid fitting curves for 1/*x* (**u**) and *DA* (**v**) time courses shown in **s–t**. All shaded regions represent SEM.

Next, we quantified system complexity by computing the dimension of activation (*DA*, **Methods**) from the LFP signals. *DA* and E/I are tightly linked because the E/I balance directly determines the net excitation a neural circuit receives. Note that while the estimation of E/I balance is a spectral approach, the computation of *DA* treats a signal as a trajectory, agnostic to any spectral information. Strikingly similar to the E/I profile, *DA* increased during Transition, peaked around EoB, and stabilized onward (**Figures 3d–e**). Results of all other individuals are shown in **Figures S1j–r**. *DA* was significantly different across stages, at both the population level (**Figure 3f**) and the individual-subject level (**Figures S2k–t**). All brain regions exhibited higher *DA* during Task Engagement than during Anesthesia (**Figure 3h**).

Taken together, results from two independent routes, one spectral and the other spectral-free, converge on the idea that the brain became more active, excitatory, and complex during the emergence of consciousness from anesthesia. The degree of excitation and activation continuously heightened after anesthesia ceased, yet stopped surging and plateaued after the appearance of behaviors, in accordance with the finding that electrophysiological change points were concentrated in Transition.

Next, we examined if different brain regions demonstrated differential activation patterns during the recovery process through the lens of E/I and *DA*. Pairwise comparisons of 1/*x* or *DA* magnitudes between all combinations of region pairs were performed using the linear mixed model (LMM, FDR-corrected, **Methods**) to account for the inconsistent channel location distribution across subjects, which is typical of human sEEG studies. Temporal and parietal structures started on equal footing—their 1/*x*and *DA* values showed no significant difference during Anesthesia, but continuously diverged during Transition, and further separated upon TE (**Figures 3m, r**). In contrast, frontal 1/*x* and *DA* were higher than those of the temporal lobe during all stages (**Figures 3k, p**). Other combinations showed slightly different results between 1/*x* and *DA*; however, the general trend remained highly consistent. These results highlight the nuances in the activation mechanism across different brain regions during the reconstitution of consciousness from anesthesia.

Both 1/*x* and *DA* of the temporal lobe were significantly lower than those of the basal ganglia, diencephalon, frontal lobe, insula, and parietal lobe (**Figures 3i–r**). The black horizontal bars in each panel mark the time points when LMM returned a significant difference (FDR-corrected *p* < 0.05) in 1/*x* or *DA* between two brain regions. Results of all other pairwise comparisons (e.g., insula vs. cingulate) are shown in **Figure S3**. Although the parietal lobe also exhibited differences in activation metrics from other regions, they were less pronounced than those seen for the temporal lobe. **Figures 3s** and **3t** juxtapose 1/*x* and *DA* traces from all regions, and **Figures 3u** and **3v** depict their sigmoid fitting curves (**Methods**). These results suggest that the temporal lobe regains consciousness differently from other brain regions.

### Pronounced alpha periodicity and synchronization during Anesthesia

So far, we have described how the brain exits anesthesia and enters consciousness at large; next, we aim to “zoom in” on the neural dynamics and identify more specific biomarkers across stages. Contrasting the power spectra during Anesthesia and TE (**Figures 4a–c**) reveals higher power at slower frequencies (delta, theta, alpha, and beta) during Anesthesia and the reverse during TE. The top and bottom horizontal bars in **Figure 4c** display the proportion of subjects whose power in a specific band during Anesthesia was greater or smaller than during TE (signed-rank test, *p* < 0.05). These results reveal a shift in dominant frequencies from anesthesia to emergence.

**Figure 4.**
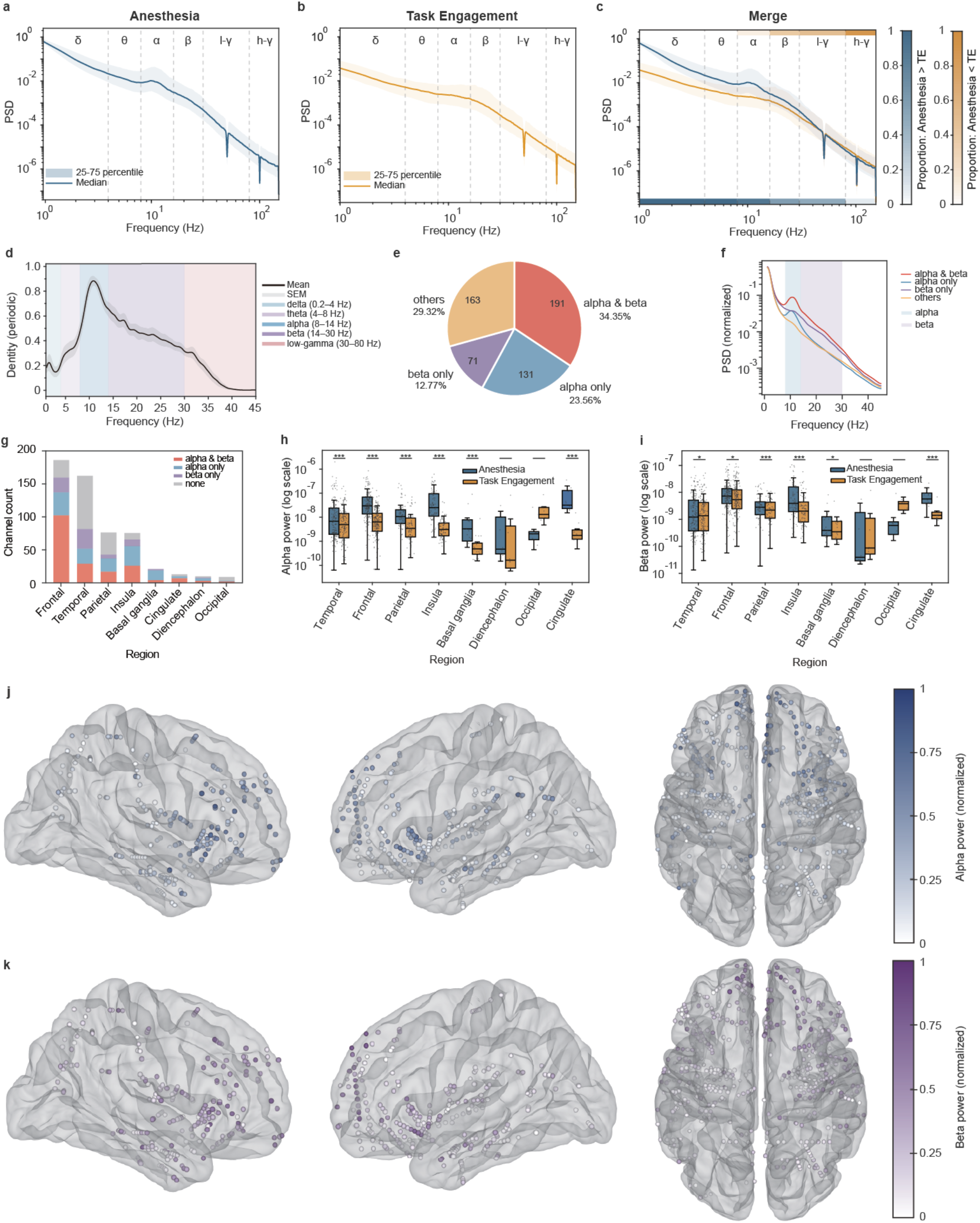
Anesthesia was marked by alpha and beta periodic activities. **a**. Power spectrum during Anesthesia. l-γ = low-gamma and h-γ = high-gamma. The line shows the median power spectral density across all subjects. Notch filters at 50 Hz and its harmonics were applied to remove AC line noise. To examine whether the power in a given frequency band differed between Anesthesia and TE within each subject, we applied the signed-rank test (α = 0.05) to compare the power values of all channels during the two stages. **b**. Power spectrum during TE in the same format as **a. c**. Merge of **a** and **b**. Top color bars indicate the percentage of subjects whose power in a specific frequency band during Anesthesia was smaller than during TE; bottom color bars vice versa. **d**. Power spectrum of reconstructed periodic components from all subjects. **e**. Percentage of channels that had significant alpha, beta, alpha + beta periodic activities, or otherwise. **f**. Median PSD of each category in **e. g**. Proportion of channels in each category, separated by brain region. **h**. Comparison of alpha power during Anesthesia and TE in each brain region. Each data point represents the median power of one channel during Anesthesia or TE. **i**. Results of beta power shown in the same format as **h. j**. Alpha power in channels showing significant alpha periodicity. Left panel: right lateral view; middle panel: left lateral view; right panel: bottom view (top: frontal; bottom: occipital). **k**. beta power in channels showing significant beta periodicity. Same view arrangement as in **j**.

A bump in the spectral density at the alpha band during Anesthesia (**Figure 4a**) implies a periodic neural oscillation at the alpha frequency that stood out from aperiodic, background activity. We leveraged spectral parameterization (SP, **Methods**) to identify the periodic components in LFP signals. Reconstructing only periodic components revealed prominent alpha and beta periodicity (**Figure 4d**). The alpha band power showed a higher, more sharply defined spectral peak, suggesting a strong and narrowband alpha oscillation. Across all channels, 57.91% revealed alpha periodicity, 34.35% exhibited both alpha and beta periodicity, and 47.12% showed beta periodicity (**Figure 4e**). For each category of channels, the median power spectral density is shown in **Figure 4f**. Channels with significant alpha/beta oscillations were widely distributed in the brain rather than confined to a few clusters (**Figures 4g, j–k**). Finer observation unpacked that both alpha and beta power demonstrated an anterior-posterior gradient (**Figures 4j–k**)—more frontal regions showing higher power, consistent with a previous study reporting alpha anteriorization^11^. **Figures 4h–i** revealed decreasing alpha and beta power from Anesthesia to TE in most regions.

Given that alpha and beta oscillations were widely distributed, it is natural to ask whether such prevalence engenders large-scale synchronized activities. Therefore, we calculated global coherence (GC, **Methods**), which incorporates both phase and power information, to quantify the strength of synchronization across different frequencies. During Anesthesia, GC peaked at 10.8 Hz in the alpha band (**Figure 5a**), meaning that neural coupling at 10.8 Hz was the strongest among all synchronous activity. There was no prominent synchronization at other frequencies and no consistent synchronization profile during TE (**Figure 5b**) or FW (**Figure 5c**). The absence of strong beta synchronization during Anesthesia might result from a much wider beta oscillation bandwidth than that observed for alpha (**Figures 4d, f**). The wider the frequency range the oscillations occupy, the less likely they are to oscillate at the same pace to become synchronized.

**Figure 5.**
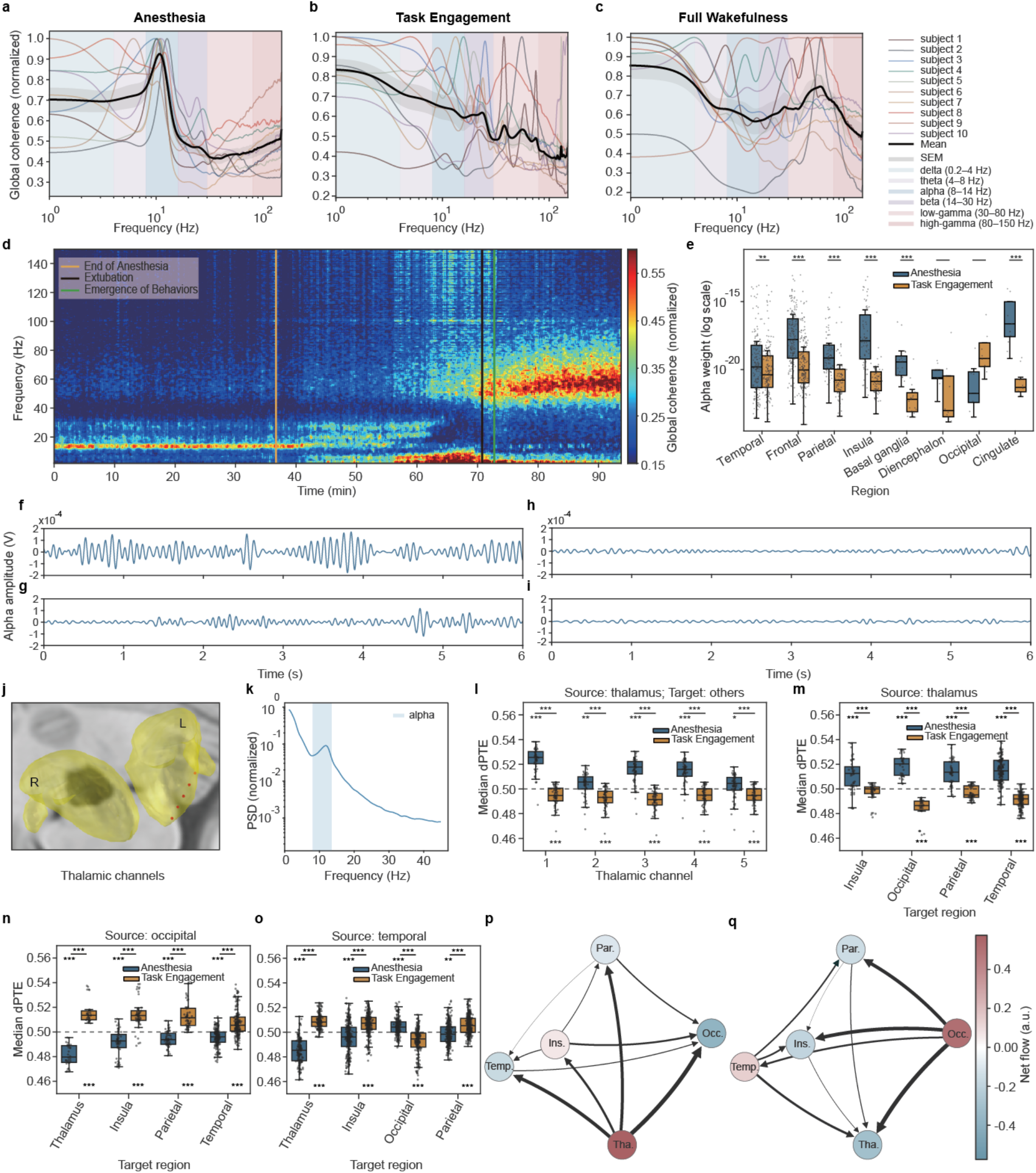
Anesthesia was marked by global alpha synchronization and directed thalamocortical information flow. **a–c**. Global coherence (GC) profiles during Anesthesia (**a**), Transition (**b**), and TE (**c**). Thicker black line shows the average GC during across all subjects. Shaded regions indicate s.e.m. Each colored line in the back shows the GC for each subject, which was determined as the median GC across time during each stage, then normalized to its minimum and maximum values. **d**. Visualization of GC across time in one subject. Higher GC indicates stronger synchronization. **e**. Contribution of alpha oscillations (alpha weight) to the global state of neural synchrony. Paired boxplots illustrate alpha weights under Anesthesia versus TE in each region (signed-rank test). **f–i**. Example raw alpha waveforms during Anesthesia (**f**), Transition (**g**), TE (**h**), and FW (**i**). **j**. Location of channels in the thalamus. **k**. Median PSD of all thalamic channels. **l**. Comparison of *dPTE* between Anesthesia and TE for every thalamic channel. The thalamus was defined as the source and other regions as targets in the *dPTE* calculation. Each data point indicates the median *dPTE* value of a source-target channel pair during a stage. Comparison with the baseline (*dPTE* = 0.5) and comparison between stages were carried out with the signed-rank test. **m**. Comparison of *dPTE* between Anesthesia and TE, combining all thalamic channels but separated by target region. **n**. Comparison of occipital (source) → others (target) *dPTE* between Anesthesia and TE. **o**. Comparison of temporal (source) → others (target) *dPTE*. **p–q**. Overview of the connectivity network during Anesthesia (**p**) and TE (**q**). Edge width indexes the magnitude of connectivity, and the arrowhead marks the direction of connectivity. Red nodes indicate inferred sources and blue nodes inferred targets (**Methods**). Node color shade reflects the net connectivity magnitude between that node and all others.

A visualization of the GC profile throughout the entire session in one subject (**Figure 5d**) reveals a strong, persistent, and sharply focused alpha synchronization during Anesthesia. Soon after anesthesia terminated, such robust alpha synchronization became diluted and almost completely disappeared shortly before Extubation / EoB, and thereafter remained subtle throughout TE. GC profiles of all other subjects are displayed in **Figure S4**. We computed the alpha weight (**Methods**) to estimate the contribution of alpha synchronization to the overall state of synchrony across different brain areas and stages. In most regions, alpha weight was significantly higher during Anesthesia than during TE (**Figure 5e**). The decoupling of alpha oscillations after EoA may arise from a decrease in amplitudes and an increase in frequency (**Figures 5f–i**). Taken together, these results suggest that alpha periodicity and alpha global synchrony are salient electrophysiological signatures of unconsciousness.

### Stage-dependent direction of information flow within the thalamocortical network

We have shown that alpha periodic activities were widely distributed during Anesthesia, which formed a robust and large-scale synchronization network peaked at 10.8 Hz. It is therefore important to know how information flows within this network to sustain the state of unconsciousness. The thalamus is thought to be highly involved in consciousness-related processes through the thalamocortical network^54^. Nevertheless, human neurophysiological data from the thalamus are extremely scarce compared with other brain regions, constraining our understanding of thalamic mechanisms in humans—especially regarding aspects of consciousness and mentation that are either unique to humans or differ substantially from those in other animals. ECoG and sEEG are the two major types of electrodes for invasively tracking seizure foci. ECoG records from the cortical surface and barely captures subcortical activity. SEEG electrodes, despite their accessibility to deep structures, are less frequently placed in the thalamus due to limited clinical indications and technical challenges. Only since the past decade has thalamic sEEG gained attention because of further understanding of thalamic involvement in epilepsy networks, more advanced surgical technology, and exploration of thalamic targets for neuromodulation^55^. One subject received sEEG implantation in the left dorsal thalamus (**Figure 5j**); therefore, we investigated the information flow between thalamus and other brain regions, given its presumptive role as a gatekeeper^14^, pacemaker^56^, and synchronizer^57^ governing distinct conscious states.

Examination of the power spectral density of thalamic channels revealed conspicuous alpha oscillation (**Figure 5k**). We investigated information flow orchestrated by alpha frequency, using directed phase transfer entropy (*dPTE*, **Methods**). *dPTE* reveals the direction and magnitude of information transfer between signals, by measuring how the phase of one signal influences or predicts the future phase of another signal. A *dPTE* value greater than 0.5 indicates directional information flow from the defined source to the defined target; a value of 0.5 indicates no net directional information; and a value less than 0.5 indicates flow from the target to the source. **Figure 5l** shows the information flow from each thalamic channel to all other simultaneously recorded channels. All subsets exhibited significant difference from the 0.5 baseline (signed-rank test, *p* < 0.05). Furthermore, all thalamic contacts demonstrated significant difference in *dPTE* between Anesthesia and TE (signed-rank test, *p* < 0.05). Anesthetic stage was marked by a strong directional phase binding from the thalamus to other regions (blue boxes), and such binding both reversed and attenuated during TE (orange boxes). **Figure 5m** showed the same data, but grouped not by the source channels but by target regions. All thalamus-target region pairs showed significant *dPTE* differences from baseline, as well as significant differences between Anesthesia and Task Engagement. Regions less than 5 channels were not included.

Next, we investigated information flow among all recording sites within this subject. The occipital lobe exhibited completely opposite *dPTE* profile to that of the thalamus (**Figure 5n**). During Anesthesia, *dPTE* from the occipital lobe to other regions was significantly lower than 0.5, indicating that the alpha oscillations transmitted toward the occipital area. This direction was reversed during TE. Therefore, while the thalamus mainly served as an information sender during Anesthesia, the occipital lobe acted as an information receiver. Similarly, the temporal lobe was another major alpha recipient, albeit not from the occipital lobe (**Figure 5o**). Other regions did not reveal strong regularity (**Figure S5**). **Figures 5p–q** show a schematic of the *dPTE*-based network, in which all regions containing ≥ 5 channels were included. During Anesthesia, the thalamus was the dominant information source, and the occipital lobe the primary target. However, their roles reversed upon TE. Although based on preliminary observations from a single subject, these results suggest a potential mode switch in the thalamocortical network during the transition from unconsciousness to consciousness, warranting further investigation and validation.

### Aperiodic slow waves during Anesthesia

Slow-wave (0.2–2 Hz) activity did not exhibit pronounced periodicity compared with alpha and beta bands (**Figure 4d**); however, it showed sustained high power throughout Anesthesia (**Figure 2a**). Multiple studies have observed strong, high-amplitude delta band (0–4 Hz) or slow-wave (SW, 0.2–2 Hz) activity during states of insensibility, such as N3 sleep, anesthesia, and coma^58^. However, the specifics of such slow waves remain nebulous. It is unclear whether they are periodic or aperiodic, and how they may change across different conscious states. Answering these questions 1) establishes a more precise slow-wave biomarker for states of unconsciousness, 2) improves interpretation of iEEG or EEG-based measures of conscious states in both clinical and research settings, and 3) propels investigation of the underlying neurobiological mechanisms that differentiate between periodic and aperiodic activities.

Not only spectral parameterization but also direct visualization of the waveform indicates that the slow wave was aperiodic during Anesthesia (**Figure 6a**). Compare its irregular, asymmetric waveform with the alpha oscillations in **Figure 5f**. The amplitude of slow-wave activity decreased after anesthesia ceased and remained low throughout TE (**Figure 6e**). Data from each individual subject are shown in **Figure S6**. The half-wave duration shortened across stages (**Figure 6f**), which corresponded to its increased frequency (**Figure 6g**). Due to SW’s asymmetric waveform, “half-wave duration” (**Methods**) is a more appropriate metric than “period”. For all brain regions, slow-wave amplitude was significantly higher during Anesthesia than during TE (**Figure 6h**). The half-wave duration showed the same downward trend in all regions except diencephalon and cingulate (**Figure 6i**). The large variance in half-wave duration during Anesthesia corroborated that slow-wave activity at this stage was aperiodic. SW frequency turned significantly faster from Anesthesia to TE in most regions, measured with the incidence of half-wave per second (**Figure 6j**). In contrast to alpha and beta oscillations, slow-wave strength did not show a salient regional gradient (**Figures 6h, k**). Collectively, slow-wave activity marks the state of unconsciousness, with an aperiodic nature that was distinct from alpha and beta periodicity. Such discrepancy in electrophysiological fingerprint might arise from distinct underlying neurobiological mechanisms—those generating stable neuronal rhythms versus those devoid of pacemaker cells or synchronized ensembles.

**Figure 6.**
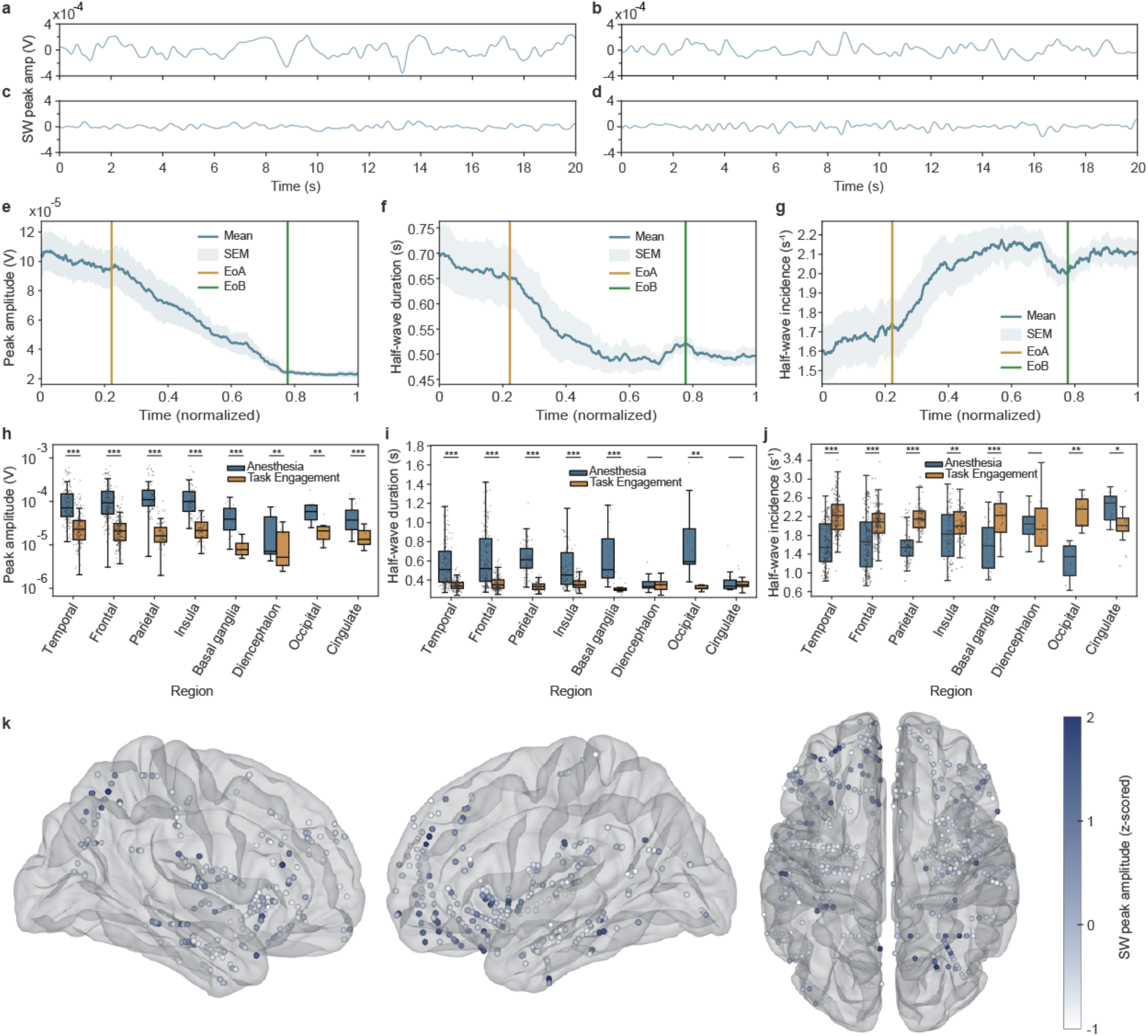
Aperiodic, high-amplitude slow waves characterized Anesthesia. **a–d**. SW waveform examples during Anesthesia (**a**), Transition (**b**), TE (**c**), and FW (**d**). **e–g**. Change in peak amplitude (**e**), half-wave duration (**f**), and half-wave incidence (**g**) of SW throughout the recording sessions, normalized onto a common time axis, and then averaged across all subjects. **h–j**. Region-wise comparison of peak amplitude (**h**), half-wave duration (**i**), and frequency (**j**) between Anesthesia and TE. **k**. Normalized slow-wave peak amplitudes of all channels in the gray matter. Left panel: right lateral view; middle panel: left lateral view; right panel: bottom view (top: frontal; bottom: occipital).

### Change in cross-frequency coupling from Anesthesia to Task Engagement

So far, we have identified neural signals in the slow-wave, alpha, and beta bands as key players during Anesthesia. The following question is how these neural activities at different frequencies interact and how such interactions change across stages. A canonical metric in LFP/EEG studies for examining the orchestration between low- and high-frequencies is the phase-amplitude coupling (PAC)^59^, specifically, the relationship between low-frequency phases and high-frequency amplitudes. We calculated SW-alpha and SW-beta PAC using the modulation index (MI) approach (**Methods**). To evaluate if the MI of a channel was significantly above chance level during Anesthesia, we applied a permutation-based phase shufling test (**Methods**). The percentages of channels exhibiting significant PAC during Anesthesia were 25.2% (SW-alpha), 21.6% (SW-beta), and 12.1% (both), among which frontal SW-alpha PAC featured most prominently (**Figures 7a–c**). Subsequent analyses in this section focused on channels with significant MI only.

**Figure 7.**
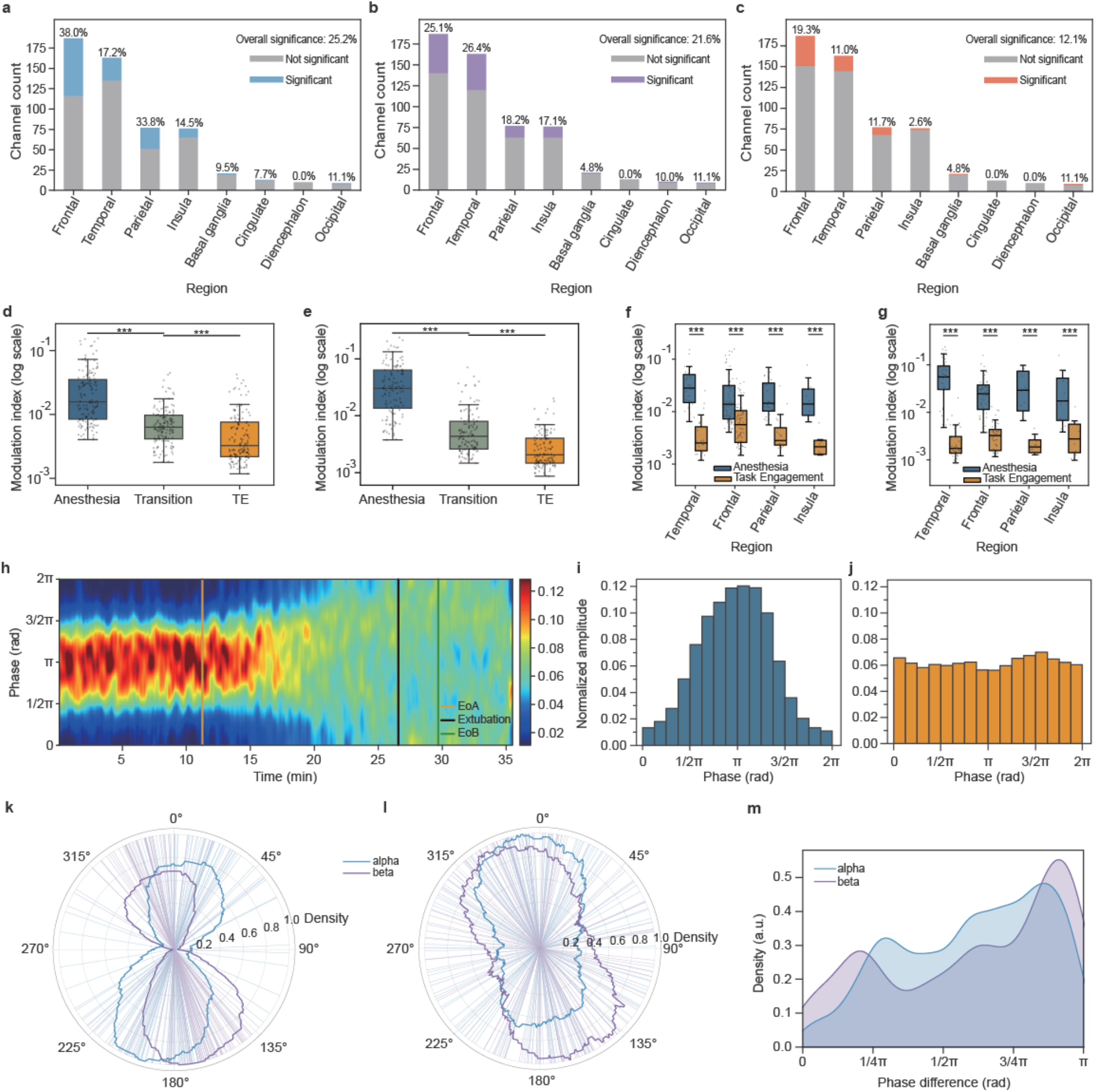
Profile of phase-amplitude coupling from unconsciousness to consciousness. **a–c**. Proportions of channels with significant SW-alpha PAC (**a**), SW-beta PAC (**b**), and both (**c**). “Overall significance” indicates the percentage of channels demonstrating significant PAC (across all channels). **d–e**. Attenuating SW-alpha (**d**) and SW-beta (**e**) PAC strength across stages. **f**. Comparison of SW-alpha PAC between Anesthesia and TE within each region. **g**. Comparison of SW-beta PAC between Anesthesia and TE within each region. **h**. Correlation between alpha amplitude and slow-wave phase in an example channel in the right middle temporal gyrus. Phases were divided into 16 bins (y axis). Within each column of time (x axis), the 16 amplitude values corresponding to the 16 bins of phases were normalized such that their sum equaled 1. **i–j**. Normalized amplitude along phase bins during Anesthesia (**i**) and TE (**j**). **k–l**. Preferred phases during Anesthesia (**k**) and TE (**l**). **m**. Kernel (Gaussian) density estimation of preferred phase difference between Anesthesia and TE. For any two phases, the shorter path along a circle was taken; therefore, the maximal phase difference was π.

Both SW-alpha and SW-beta PAC demonstrated successive declines across stages (**Figures 7d–e**). **Figure 7f** shows the comparison of SW-alpha PAC between Anesthesia and TE in each brain region. Regions with fewer than 5 channels of significant MI were not shown. All remaining regions demonstrated remarkable PAC differences between the two stages. Same patterns were observed for SW-beta PAC (**Figure 7g**).

Previously, we computed MI to measure how strongly the amplitude of high-frequency activity is coupled to the phase of low-frequency activity. Next, we examined what specific phases orchestrated the observed PAC and whether these critical phases shifted across stages. **Figure 7h** presents a visualization of the coupling between alpha amplitudes and slow-wave phases during the entire recording session from one example channel. During Anesthesia, higher alpha amplitudes tended to fall around π, while such preference started to dissipate in the middle of Transition (see also **Figures 7i–j**). Population results indicated that during Anesthesia, the preferred phases formed two clusters, a larger one around π and a smaller one around zero (**Figure 7k**), both of which subsequently dispersed during TE (**Figure 7l**). **Figure 7m** depicts the distribution of preferred phase shift from Anesthesia to TE. These results suggest that the binding of slow wave and alpha/beta activities through phase-amplitude coupling may sustain the state of unconsciousness, whereas the phase-amplitude decoupling foretells the emergence of consciousness.

### Differential recovery trajectories of behavioral performance across tasks and individuals

Cognitive tasks were administered repeatedly throughout the entire recording procedure to instantaneously assess subjects’ states of consciousness. During Anesthesia, there was no overt response or behavior. After EoA, subjects started to wake up and regain awareness to self and surroundings. Brain states incrementally resembled their modes during FW, such that conscious behaviors could emerge. Specific task commands and the functions they measure are delineated in **Figure 1b**. Each block contained four trials that tested Movement to Command (MC), Verbal Communication (VC), Visual Recognition (VR), and Functional Object Use (FO) respectively. Trials requiring visual functions (VR and FO) were given only after eye opening. The number of blocks completed by each subject is shown in **Table S3**.

On average, 19.8 ± 8.7 (mean ± SD) minutes elapsed from EoA to EoB. Subjects performed the task increasingly well after EoB and subsequently throughout TE (**Figure 8**). **Figures 8a–d** record the behavioral performance of all subjects across four tasks. An accuracy of 0 equated to no behavioral response; 0.5 equated to incomplete or incorrect responses (e.g., a partially completed motor movement or an incorrectly stated birthday); 1 equated to perfect performance. Visual Recognition (**Figure 8c**) displayed a more tortuous recovery process than the other three functions. To model behavioral dynamics at population level, we employed a Bayesian state-space model (**Methods**) to estimate the time-varying probability and credible interval for performance scores (**Figures 8e–i**). Temporal normalization was implemented such that t = 0 indicates EoA, and t = 1 marks EoB. The expected score at t = 1 served as an estimate of the initial performance. MC showed the highest initial expected score and VR the lowest, and such relative magnitudes persisted until near-perfect performance had been achieved for all tasks (**Figure 8i**). Therefore, simple movement to command, which demanded basic auditory, comprehension, and motor functions, came back first, followed by the reconstitution of vision that was more challenging to recover. The differential recovery trajectories were not due to any difference in task difficulty, given the perfect performance during Stage 4 (**Figure S7**).

**Figure 8.**
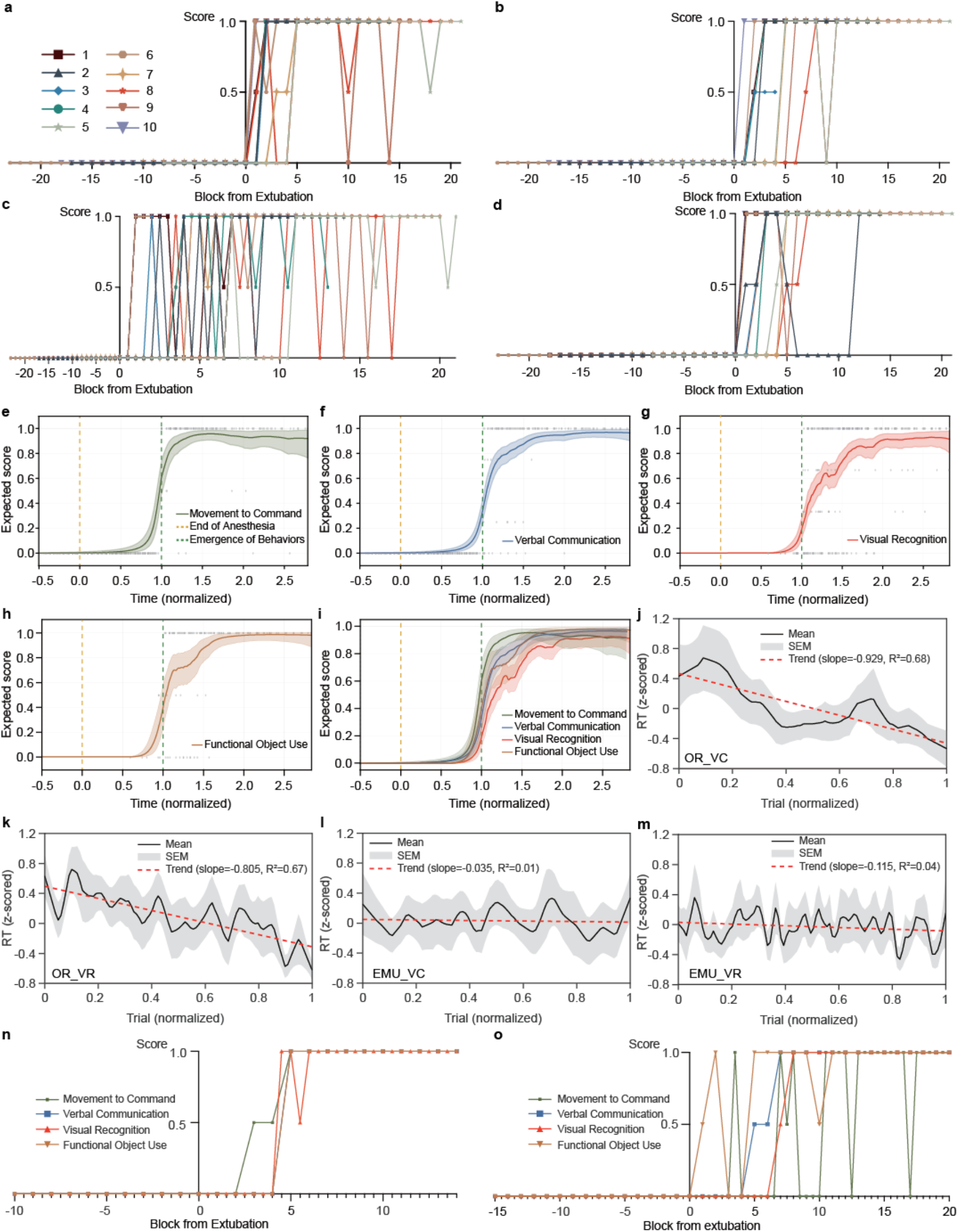
Behavioral performance. **a**. Subject performances of MC. Each colored line represents one subject. **b–d**. Performances of VC, VR, and FO, in the same format as **a. e–h**. Expected scores of MC, VC, VR, and FO, on a normalized time axis, estimated from a Bayesian state-space model. **i**. Overlay of panels **e–h. j**. Population-level reaction time trajectory of the VC in the OR. For each subject, each trial returned one RT value. To display group average, the z-scored RT vectors from all subjects were linearly interpolated to a common normalized trial axis. The red dashed line indicates a least-squares linear regression fitted to the group-mean trajectory. **k**. RT of the Visual Recognition task in the OR. **l**. RT of VC in the epilepsy monitoring unit when wide awake. **m**. RT of VR in the EMU. **n**. Example task performance of one subject with fast recovery. **o**. Example task performance of one subject with slow, uneven recovery.

Another behavioral indicator of the recovery of consciousness was the decrease in reaction time (RT). Verbal responses for VC and VR were analyzed offline to determine the RTs (**Methods**). In the OR, RT significantly shortened across trials (**Figures 8j–k**), reflecting a gradual enhancement of the level of consciousness. This was unlikely due to a learning effect given that patients were fully familiarized with the task and practiced it before the surgery. Furthermore, no significant decrease in RT was observed during FW (**Figures 8l–m**). Thus, the reduced reaction time and improved accuracy on these simple exams—which tap into the most fundamental functions of everyday life—bespeak an ongoing reconstruction of consciousness after its pharmacological deconstruction.

So far, we have demonstrated that different functions recuperated not simultaneously but by order. We next sought to investigate if there was subject-wise difference in the rate of recovery. **Figure 8n** shows the task performance from one subject with a fast return of conscious behaviors in all tests. In contrast, **Figure 8o** displays a slow and difficult progression from another subject with a prolonged period of fluctuation in the performance score. To test whether there was significant individual variability considering all subjects, we used the “number of blocks (*t*) to consistently perfect performance” as an index for the rate of recovery in each task (**Methods**). Parametric bootstrapping on *t* revealed significant subject-wise heterogeneity in VC (*Var*(*t*) = 12, *p* = 0.001), VR (*Var*(*t*) = 13.44, *p* = 0.038), and FO (*Var*(*t*) = 11.28, *p* = 0.042). Only MC showed no evidence for inter-subject difference (*Var*(*t*) = 2.86, *p* = 0.921). Therefore, the restoration of conscious function unfolded at different paces across individuals, suggesting an idiosyncratic nature of the recovering brain.

### Ceding power to high-gamma and freeing consciousness

We predicted that certain channels would exhibit continuous changes in their electrophysiological properties from anesthesia to emergence, gradually converging toward the neural correlates of conscious behaviors observed during FW. Two lines of evidence let us narrow down the NCC candidates to the high-gamma power. First, high-gamma activity is a cosmic indicator of assorted cognitive processes, which have been demonstrated by an avalanche of research^60–62^. Second, contrasting the power spectra across stages revealed that only the high-gamma band exhibited higher power during Stages 3/4 than Stages 1/2 in the majority of subjects (**Figure 4c** and **Figure 9a**). Indeed, high-gamma power effectively captured distinct types of selectivity during FW (**Methods**). In addition to neural responses sensitive to each task, we incorporated modality selectivity, including auditory and verbal selectivity into the analysis. Auditory responses pertained to all tasks because audio commands were provided. Verbal selectivity applied to VC and VR, where spoken answers were required. For this section only, we examined both gray and white matter channels to enlarge the otherwise small sample size, inasmuch as channels in the white matter could demonstrate salient task or modality selectivity (**Figures S8–10**). Previous research has suggested that white matter LFPs can be biologically meaningful, reflecting signals from nearby or distant gray matter^63,64^. **Figures 9b–f** and **Figures S11b, d, f, h** display high-gamma activity during FW from example channels showing auditory, verbal, MC, VR, and FO selectivity respectively. VC-selective channels were not considered because they were scarce in number and weak in response. All task- and modality-selective signals are laid out in **Figures S8–10**.

**Figure 9.**
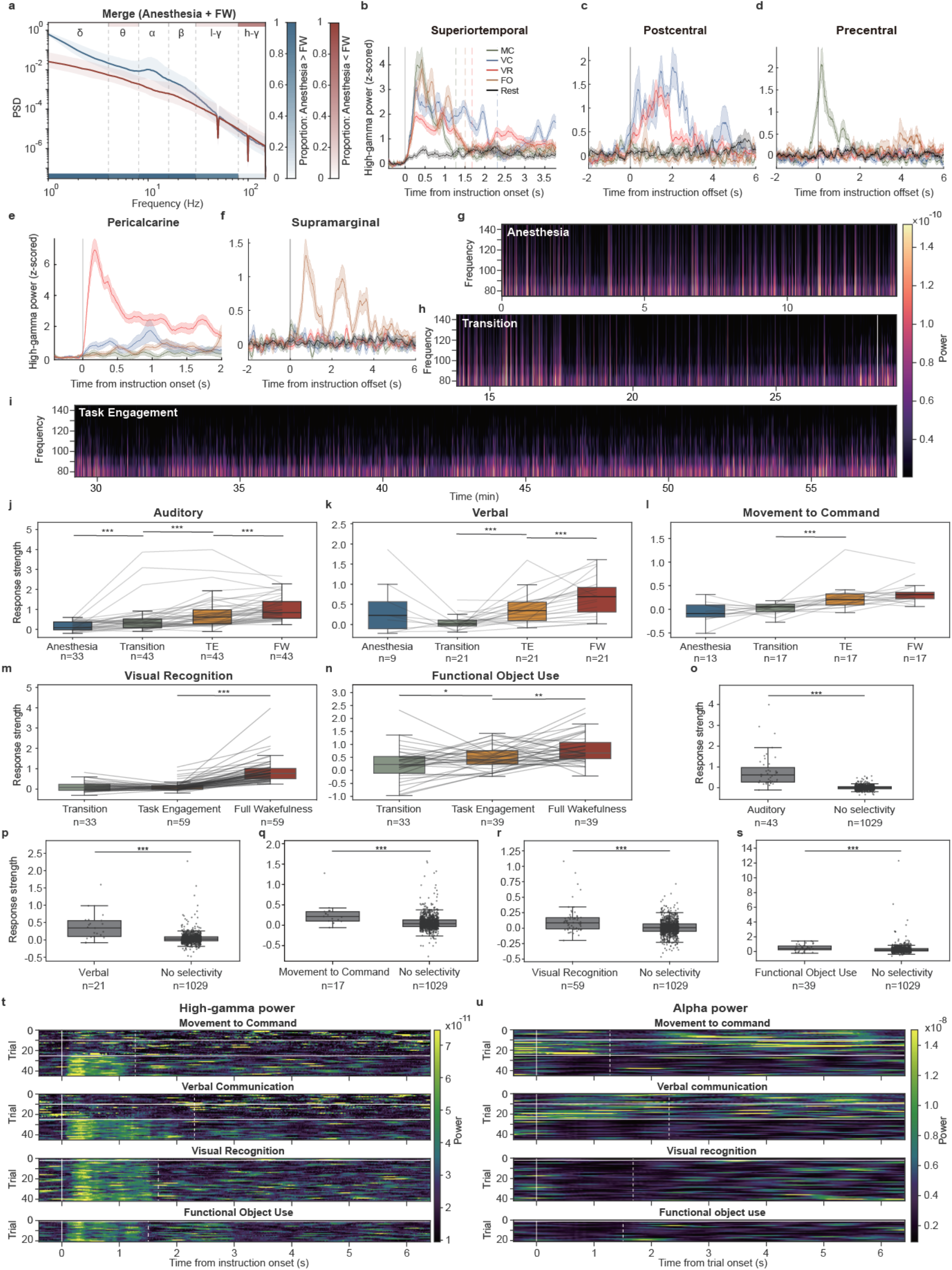
High-gamma activity supports conscious behaviors. **a**. Merged power spectra of Anesthesia and FW. Top color bars indicate the percentage of subjects whose power in a specific frequency band during Anesthesia was smaller than during FW; bottom color bars vice versa. **b**. Example auditory-selective channel, with sustained high-gamma activity when audio was on. Signals were aligned to instruction onsets. Dashed lines show the offsets of audio commands. No high-gamma activity was observed during resting periods. **c**. Example verbal-selective channel, with higher high-gamma power after instruction offset, representing neural correlates of verbal responses for VC and VR. Signals were aligned to instruction offsets. **d–f**. Example MC-selective (**d**), VR-selective (**e**), and FO-selective channels (**f**). **g–i**. Appearance of high-gamma power during Anesthesia (**g**), Transition (**h**), and FW (**i**) from an example channel. The vertical white line in **h** represents the timing of Extubation. **j–n**. Comparison of response strength (**Methods**) for channels demonstrating selectivity across stages. Each line represents data from a single channel, connecting the median response strength across all trials in each stage. Pairwise comparisons were conducted using the one-tailed Wilcoxon signed-rank test. **o–s**. Comparison of response strength during TE between channels showing each type of selectivity and channels showing no selectivity at all. Comparisons were conducted using the two-tailed Wilcoxon rank-sum test. **t**. Raster plot displaying the high-gamma power in every trial in the OR from an auditory-selective channel. Vertical solid lines indicate instruction onsets. Dashed lines mark instruction offsets. Each small panel may contain two (EoA then EoB) or one (EoB) horizontal white line, depending on the number of blocks completed in each stage and whether the task was administered only after eye opening (VR, FO). **u**. Per-trial alpha power in the same format as in **t**.

First, the morphology of high-gamma activity underwent a profound transformation (**Figures 9g–i**). Anesthesia and Transition were characterized by stochastic bursts of power, while TE showed more sustained and stable activity. Such distinction is even more appreciable in the raster plots displaying the high-gamma dynamics in every trial (**Figure 7t** and **Figures S11a, c, e, g**). **Figure 7t** plots the high-gamma responses during all MC trials in the OR, from an auditory-selective channel. High-gamma activity during Anesthesia and Transition revealed no consistent structure. In stark contrast, during TE, the high-gamma response was strictly stimulus-locked, with stronger power sandwiched between the onset and offset of the audio command. Such change was rather sudden than gradual. As soon as conscious behaviors took shape, the mode of high-gamma activity became orderly and predictable from trial to trial. Other types of selectivity demonstrated similar effects (**Figures S11a, c, e, g**). Next, examining the raster plots in the OR and EMU revealed that the high-gamma morphology was far more congruent between TE and FW than between Transition and TE (**Figure S11**).

Second, the perfection of conscious behaviors was accompanied by an increase in high-gamma activity. **Figures 9j–n** depict how high-gamma response strength (**Methods**) changed across stages. Auditory responses steadily rose from Anesthesia to FW in a stepwise manner (**Figure 9j**). All types of selectivity except VR (**Figure 9m**) showed a significant increase in response strength from Transition to TE. This exception is consistent with the previously presented behavioral result showing that vision was the most difficult function to recover. In a similar vein, all types of selectivity except MC (**Figure 9l**) showed a significant increase in response strength from TE to FW. This suggests that MC recuperated most quickly and fully, as the response strength during TE was already on par with that during FW. No data were available during Anesthesia and during portions of the Transition for VR and FO, as these tasks were administered only after eye opening (**Figure 1b**). Channels without any type of selectivity exhibited significantly lower response strengths than those observed from selective ones during TE (**Figures 9o–s**), suggesting that the salient increase in high-gamma activity was not universal, but dictated by current demands. The alpha band showed the opposite effect—higher power during Anesthesia and Transition but minimal activity during TE (**Figure 9u**)—consistent with the earlier finding that strong alpha periodic activity dominated Anesthesia and then attenuated as consciousness revived.

In short, the neural circumstance attending the moment of the emergence of conscious behavior was a dramatic metamorphosis in the high-gamma activity. From unconscious to conscious states, from zero to perfect behaviors, high-gamma power steadily increased, in those sites that are responsible for various aspects of cognition. The primary neural correlate of the emergence of conscious behaviors is activity in the high-gamma band.

## Discussion

How consciousness emerges is one of the most profound questions in modern science. In this work, we provide initial steps toward addressing this topic by understanding how the human brain recovers from general anesthesia, a process during which consciousness is extinguished and then revived. Specifically, we examined how brain states evolved from propofol-induced general anesthesia to emergence, characterizing the neural correlates of unconsciousness^1^, the neural correlates of consciousness^2^, and the transition between them. Multimodal cognitive scaling was administered throughout the entire procedure, titrating the level of conscious behaviors and their neural underpinnings.

States of unconsciousness or diminished conscious access are often characterized by reduced or disrupted functional connectivity among critical nodes in the conscious processing system^22,65–67^. We argue that the unconscious state is not simply a residual consequence of subtracting NCCs from the collective neural activities supporting consciousness, but is instead actively maintained by specific mechanisms that confine neuronal communication and thereby fetter conscious processing. During the reconstruction from anesthesia to emergence, certain neural processes decline while others thrive.

A central feature of this active reconstitution was the brain’s passage through a state of substantial changes and instability (**Figure 2**). Beneath this apparent chaos, however, lies structured dynamics. Power fluctuations increased across frequencies, especially in lower-frequency activity, demonstrating a stable-unstable-stable progression from Anesthesia to Task Engagement. In parallel, the brain became progressively more activated and complex (**Figure 3**) while escaping a constrained anesthetic regime and regaining the capacity for flexible, high-dimensional activity. The plateau of both E/I balance and activation dimension after EoB suggests that the return of consciousness may involve crossing a critical threshold rather than simply accumulating activation. In this sense, Transition is not a gradual amplification of the same neural processes, but a biological corridor connecting qualitatively distinct brain states.

Anesthesia was dominated by electrophysiological activity in lower-frequencies, including periodic oscillations in alpha and beta bands, as well as aperiodic fluctuations of slow wave. These activities were bound through phase-amplitude coupling, where higher alpha and beta power tended to reside at certain slow-wave phases. Besides, the strong and widespread alpha periodicity formed a large-scale synchronization peaking at 10.8 Hz. Together, these neural phenomena constitute the NCUCs associated with propofol-induced general anesthesia and may functionally suppress conscious processing.

The distinction between periodic and aperiodic activities is important because they may arise from fundamentally different neurobiological mechanisms. Simply using the phrase “frequency band” would obscure this distinction. Periodic alpha oscillations may arise from rhythm-generating neuronal ensembles^68,69^, whereas aperiodic slow waves may reflect irregular transitions between cortical UP and DOWN states^70,71^, or broader constraints on neural responsiveness^27,72^. Their common association with unconsciousness does not imply a common origin. Supporting this view, alpha and beta activity exhibited anteriorization, whereas slow waves did not. Therefore, although alpha, beta, and slow-wave activity were all prominent during anesthesia, they may arise from distinct neurobiological mechanisms and serve different functions. A precise NCUC should therefore not be defined by frequency alone, but by the joint profile of frequency, periodicity, spatial distribution, synchronization, functional coupling, and potentially other relevant dimensions not examined here.

The phase-amplitude coupling results further suggest that anesthetic unconsciousness may depend on functional interactions among distinct low-frequency motifs. During anesthesia, slow-wave phase modulated alpha and beta amplitudes, binding slow and faster rhythms into a coherent neural machinery. Through this binding, the anesthetized brain may circumscribe higher-frequency activity to specific temporal windows, thereby limiting the independence and flexibility of local neural assemblies. The dissolution of PAC may release neural populations from slow-wave control, allowing greater degrees of freedom in local neuronal actions. In this framework, consciousness emerges not simply from a reduction in low-frequency power, but from the progressive fragmentation of a temporal architecture that interlocks multiple processes to sustain unconsciousness.

The results of thalamocortical information flow provide a preliminary but provocative window into how the low-frequency regime may be coordinated. Alpha-band information flow during Anesthesia was dominated by thalamic output to cortical and subcortical targets, whereas this relationship reversed during TE. The occipital lobe showed the opposite profile, acting as a major recipient of alpha-band input during Anesthesia and shifting toward a source-like role during TE. Although these observations are based on a single subject and require validation, they raise the possibility that the thalamus may act as a state-dependent gatekeeper for conscious processing via a dynamic mode switch. This finding is particularly intriguing given that human thalamic intracranial recordings are rare, and that the thalamus has long been central to theories of arousal and consciousness^10,73,74^, as well as a primary target for neuromodulatory treatments for disorders of consciousness^75,76^.

Anatomically, the recovery of consciousness was spatially heterogeneous. The temporal lobe exhibited lower E/I balance and dimension of activation across much of the recording period. The frontal lobe showed stronger alpha and beta power during Anesthesia, implying a more inhibited or rhythmically constrained frontal state. The occipital lobe, despite limited sampling and reduced statistical power, frequently exhibited effects opposite to those observed in other regions (**Figures 4h–I, 5e, 5m–q**). These anatomical differences argue against a model in which consciousness is globally lost and globally restored in a uniform fashion, and against a model in which un/consciousness resides in a single islet.

At a pivotal moment, high-gamma activity crystallized and behavioral responses manifested. On average, 20 minutes elapsed between anesthesia ceased and behaviors took shape. High-gamma morphology changed qualitatively at EoB. During Anesthesia and Transition, high-gamma activity appeared as stochastic bursts without discernable task structure. Once behavior surfaced, high-gamma responses became orderly, event-locked, task-selective, and increasingly similar to those recorded during FW. Importantly, such high-gamma activity was distributed but not global (**Figures 9o–s, Figures S8–10**). These features distinguish patterned high-gamma activity from nonspecific activation and nominate it as a strong candidate NCC for behaviorally accessible conscious contents.

Although this study was not primarily designed to test against consciousness theories, several findings were highly germane to the instantiation of the global neuronal workspace theory (GNWT)^13,22^. GNWT posits that conscious access arises from “ignition”: a sudden, large-scale, and recurrent pattern of neuronal activation that enables information to become globally available across the brain. Although ignition classically describes how sensory information enters consciousness during wakefulness, the emergence of behaviors and its neural dynamics during recovery from anesthesia appeared highly “ignition-like”. At the time of EoB, high-gamma activity exhibited a sudden and dramatic metamorphosis, shifting from irregular bursts to organized patterns temporally locked to stimuli or responses. Although the overall high-gamma power differed across stages, no salient progressive increase could be seen within Transition or within TE. This is reminiscent of the nonlinear divergence between neural processes underlying conscious versus unconscious perception^22,77,78^. Thus, the emergence of behavioral responsiveness from anesthesia-induced unconsciousness may reflect a threshold-like cortical mobilization that resonates with the concept of ignition.

The GNW is formulated within a distributed, non-localizationist framework^22^. However, several brain regions, including but not limited to the PFC, parietal cortex, and thalamus, have received greater attention than other regions^2,22,79–81^. Throughout the history of neuroscience, the pendulum has swung back and forth between localization of function and mass action^82–84^—later reframed as distributed network^85,86^. We argue that many brain functions arise from a delicate interplay between distributed processing and localized functional hubs, and that consciousness is unlikely to differ. From unconsciousness to consciousness, some neural processes were widely distributed, including slow waves, SW-αβ PAC, and high-gamma activity, whereas others appeared more localized, such as the relatively less activated temporal lobe, the alpha/beta-dominant frontal cortex, and the mode-switching thalamic nexus. These findings help constrain consciousness theories and computational models that attempt to incorporate the neuroanatomical basis of consciousness, guide neuromodulatory strategies for treating disorders of consciousness, and provide roadmaps for basic neurobiological studies at single-neuron and circuit levels.

Although the neural substrates identified under anesthesia are by possibility specific to the pharmacological agent used, this limitation inevitably applies to nearly all forms of induced or pathological unconsciousness. For instance, neural processes identified in ketamine-maintained anesthesia may be specific to ketamine, whereas those observed in disorders of consciousness may reflect the underlying pathophysiology. More studies directly interrogating consciousness are required to disentangle candidate neural signatures from potential confounding factors. Those consistently identified across various causes of unconsciousness and consciousness may therefore represent the strongest candidates for bona fide NCUCs and NCCs. Furthermore, theories of consciousness should have explanatory power over different conscious states^22^, accounting not only for intact conscious experience but also for cases in which the conscious mind is impaired, distorted, or extinguished.

In summary, we presented a comprehensive and descriptive account of how the brain progresses from the darkness of unconsciousness to the renaissance of mentation. This process was examined at a mesoscopic level, balancing the breadth and depth of analyses and discussion. In light of that, each neural correlate identified here can be further dissected from more microscopic perspectives, and can as well be readily integrated and abstracted into higher-level theoretical frameworks. To our knowledge, no previous study has combined this level of intracranial spatiotemporal resolution and anatomical coverage, rich electrophysiological vocabulary, and multimodal task design across the continuous emergence from propofol-induced unconsciousness. Together, these findings provide a systems-level account of how the machinery of consciousness is dismantled, reconfigured, and restored.

## Methods

### Intracranial recordings, anesthesia, and epilepsy patients

Intracranial recordings of local field potentials with stereotactic electrical-encephalography (sEEG) were carried out with pharmacoresistant epilepsy patients for localizing epileptogenic foci. SEEG electrodes were from Sinovation (Beijing) Medical Technology Co., Ltd., with a contact length of 2 mm, a diameter of 0.8 mm, and an inter-contact distance of 5 mm or 10 mm (**Figure 1c**). Intracranial EEG was recorded using EEG monitoring systems from Neuracle (Shanghai, China) or Beijing Yunsound Technology Company Co., Ltd. (Beijing, China) in the operating room, and Natus (WI, USA) or Nihon Kohden (Tokyo, Japan) in the epilepsy monitoring unit. In the OR, all patients underwent propofol-induced anesthesia with laryngeal mask airway. Anesthetic depth was monitored using the bispectral index (BIS), which was maintained between 40 and 50 during Stage 1. All patients (**Table S3**) were recorded at Shenzhen Second People’s Hospital (Shenzhen, China). Consents from all patients were obtained. All procedures were approved by the hospital’s Institutional Review Board and carried out in conformity with the Declaration of Helsinki.

### Experimental details

The task paradigm was adapted from the JFK Coma Recovery Scale (CRS-R)^87^, with substantial simplification and modification of the auditory, visual, motor, oromotor/verbal, and communication function scales to accommodate patients emerging from anesthesia. One day before surgery, patients completed several practice blocks to ensure familiarity with the task. Task instructions were played through an external speaker (Newmine, Beijing, China) positioned near each patient’s head and along the body midline. Verbal responses were recorded using a Yeti microphone (Logitech, Lausanne, Switzerland) with a sampling rate of 8,192 Hz. Task images were displayed on an external monitor measuring 17.13 × 9.29 inches, hung in parallel to each patient’s coronal plane and along the midline. Images for the visual recognition task were selected from the Microsoft COCO 2017 training dataset^88^ and screened by three researchers to ensure immediate recognizability. Image categories included animal, food, indoor, and outdoor scenes, which were counterbalanced across blocks. A paper cup was placed in contact with each patient’s dominant hand, a procedure with which patients had been familiarized during the practice session. The task was programmed and presented through the Psychophysics Toolbox 3.0.19 extensions^89,90^ in Matlab_R2024b (Mathworks, Natick, MA, USA)^91^ with a MacBook Pro 2023 (Apple, Cupertino, CA, USA).

### Anatomical locations

SEEG channels were localized using the iELVis^92^ toolbox. We used FreeSurfer^93^ to segment the preimplant magnetic resonance (MR) images, upon which post-implant CT was co-registered. Channels were marked in the co-registered space using Bioimage Suite^94^. Channel locations were then determined by FreeSurfer’s volumetric brain segmentation. Cortical channels were labeled using the Desikan-Killiany^95^ atlas, and subcortical ones were labeled with the ASEG atlas^96^. Subsequently, these fine-grained locations were grouped into 8 general regions shown in **Tables S1–2**. For electrodes in the white matter, two researchers independently determined their nearest gray matter locations by visual inspection. Inconsistencies, typically due to similar distances to two gray matter regions, were resolved by a third investigator. Out of 1284 bipolarly referenced channels in total, we included 556 electrodes in the gray matter (**Table S1**) and 634 in the white matter (**Table S2**). Electrodes in the white matter were only analyzed in the last section of Results. 94 electrodes were not considered for analyses due to irrelevant locations (e.g., ventricles) or significant noise. Electrode locations were mapped onto the MNI305 average brain via affine transformation^97^ for visualization. Electrodes in the thalamus (**Figure 8a**) were displayed using the LeadDBS software^98^, which is specialized for localizing and visualizing contacts within or near deep brain nuclei.

### Preprocessing of local field potential data

All raw signals were re-referenced using bipolar subtraction. Notch filters at 50 Hz and its harmonics with a bandwidth of 2 Hz were applied to remove the AC line noise. To standardize the sampling frequency across recordings, all data were resampled to 500 Hz. Finally, a high-pass filter at 0.2 Hz was applied to remove slow signal drifts. Unless otherwise noted, all analyses were performed on these preprocessed signals.

### Time-frequency decomposition

We computed Fourier coefficients 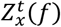 at frequency *f* for each time segment *t* from channel *x* using the multitaper method:

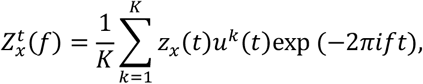

where *K* is the number of tapers, *z*_*x*_(*t*) represents the filtered signal of channel *x* at time segment *t*, and *u*^*k*^(*t*) represents the *k*-th Slepian taper. We used the MNE-Python toolbox^99,100^ to implement the multitaper method with the following parameters: n_cycles = *f*/2 (i.e., *t* length = 0.5 s), decimate = 250 (i.e., *t* step = 0.5 s), time bandwidth = 2 (i.e., *K* = 3). Spectral analyses were performed from 1 to 150 Hz in 0.5 Hz increments. Unless otherwise specified, all computational pipelines that included time-frequency decomposition adopted this multitaper method.

### Change point detection

We applied the multitaper method to obtain the time–frequency representation for each channel, which was then transformed into time-varying band power across standard frequency bands: delta (0.2–4 Hz), theta (4–8 Hz), alpha (8–14 Hz), beta (14–30 Hz), low gamma (30–80 Hz), and high gamma (80–150 Hz). To detect abrupt and significant changes in these multivariate signals, we applied radial basis function (RBF) kernel-based change-point detection. This procedure was implemented using the Python library *Ruptures*^101^, setting model=“rbf”, pen=5 for subject 4 or pen=15 for all other subjects.

### Excitatory-to-inhibitory (E/I) balance

In many neurophysiological signals such as scalp EEG and LFP, the power spectrum (*f*) exhibits a power-law decay with frequency *f*:

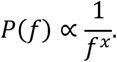

For each channel, we employed the specparam toolbox (formerly FOOOF^53^) to estimate the exponent *x* from PSDs, using a frequency range of 0.2–45 Hz, a window length of 500 ms, and a step size of 500 ms. The exponent *x* characterizes the rate at which power decreases with frequency. Previous studies^53^ have suggested that a higher *x* corresponds to a steeper spectral decay, which is associated with a lower E/I balance in neural activity. Therefore, we used 1/*x* as a proxy measure of E/I balance.

### Dimension of activation (DA)

We calculated *DA* for each channel following its standard procedures in previous research^102^. Signals were down-sampled to 256 Hz and the time delay for “delay time embedding” was fixed at 4 resampled units (15.6 ms), with the embedding dimension set to 20. *DA* was estimated as the slope of linear fit between *log*[*C*(*r*)] and *log* (*r*), in which (*r*) is the correlation sum, defined as the proportion of embedded point pairs with Euclidean distances less than or equal to *r*. For each time window, *r* was sampled as 50 logarithmically spaced values between the minimum and maximum pairwise distances, in which the linear fit was performed over the middle 60% of the *r* values. The time window was set to 15.6 s with a step size of 1.56 s.

### Region-wise comparison of E/I and DA

To compare the activation degrees among regions, we rescaled 1/*x* and *DA* time course of each channel to a normalized time axis. For a given pair of regions, only simultaneously recorded channels were taken into account (subjects who had channels in both regions). As an example, for frontal-temporal comparison, subjects who had only frontal channels and subjects who had only temporal channels were not considered. The occipital lobe was removed from this part of analysis due to the small sample size of occipital channels and the large variance in the signals. Examining 1/*x* and *DA* across regions unfolds several key activation patterns across stages. We leveraged linear mixed model for the comparison of activation metrics between regions to balance the non-independence among channels within subjects, with “region” as a fixed effect and “subject” as a random intercept. *P* values were corrected by the Benjamini-Hochberg procedure to control the false discovery rate (FDR) within each region pair. To show the overall trend in 1/*x* or *DA* for each region, we first averaged across subjects their median 1/*x* or *DA* traces, and fit the resulting time series using a sigmoid function:

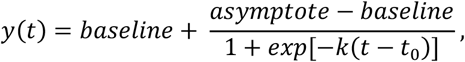

where *k* stands for the slope, and *t*_0_ stands for the inflection point.

### Spectral parameterization

For each channel, we computed stage-specific PSDs using Welch’s method in MNE-Python^99^ with a Hamming window, an FFT length of 2048 samples, no window overlap, and averaging across all segments. We employed the specparam toolbox^53^ to decompose the PSD into aperiodic and periodic components over a frequency range of 0.2–45 Hz, a maximum of 10 peaks, peak-width limits of 1–12 Hz, and a peak threshold of 1 standard deviation. The PSD is then expressed as:

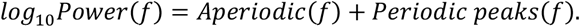

where *f* is the frequency. The aperiodic component is modeled by:

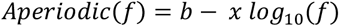

where *b* is the broadband offset, and *x* is the spectral exponent. Periodic peaks were modeled as Gaussian functions above the aperiodic PSD background. Each individual peak *n* is modeled as:

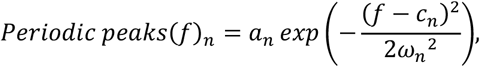

in which *a*_*n*_ stands for the peak amplitude relative to the aperiodic component, *c*_*n*_ is the center frequency, and *ω*_*n*_ is the Gaussian bandwidth parameter (standard deviation).

We used the Gaussian-modeled periodic peaks to reconstruct a normalized periodic peak distribution for each subject, and computed the mean ± SEM across subjects.

### Global coherence

The global coherence computation was adapted from previous studies^11,103^. For each time window *T* (30 s, step = 3 s), we constructed an *N* × *N* cross-spectral matrix (*f*), where the entry at row *x* and column *y* reflects the cross-spectrum between channels *x* and *y*:

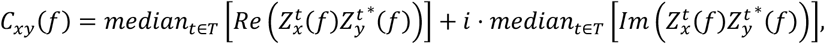

where 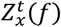 represents the Fourier coefficients at frequency *f* for each time segment *t* from channel *x*, and 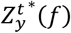 is the complex conjugate of the Fourier coefficients at frequency *f* for each time segment *t* from channel *y*. The eigendecomposition was then applied to the cross-spectral matrix *C*(*f*):

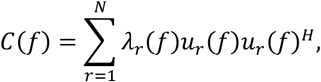

where *N* is the number of channels, *λ*_*r*_(*f*) denotes the *r*-th eigenvalue of *C*(*f*), and *u*_*r*_(*f*) denotes the eigenvector associated with *λ*_*r*_(*f*). Assuming the eigenvalues *λ*(*f*) are sorted in descending order, global coherence at frequency *f* is defined as:

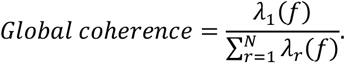

To perform a low-rank approximation (LRA) of (*f*), we retained only the largest eigenvalue and its corresponding eigenvector:

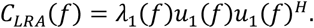

From the low-rank approximation of the cross-spectral matrix, the channel-specific weight at frequency *f* for channel *x* is defined as:

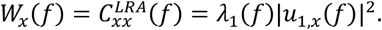

In our analysis, we estimated the alpha weight for each channel using the median weights across the frequency range *F*_*α*_ from 8 Hz to 14 Hz and time *T*_*s*_ in each stage:

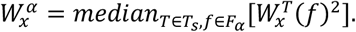

### Directed phase transfer entropy (dPDE)

We computed the *PTE* between two signals following its standard procedures^14,104^. The *PTE* between two phase series *θ*_*x*_ and *θ*_*y*_ was calculated as:

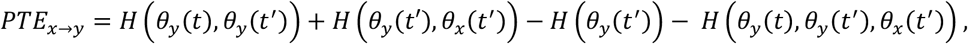

where *θ*_*x*_(*t*^’^) and *θ*_*y*_(*t*^’^) are phase states at previous time point *t*^’^ = *t* − *δ*, and the lag *δ* was set at 20 ms. The entropy *H*(*u*) was estimated using histogram-based empirical probability distributions:

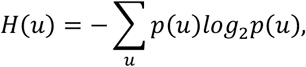

where *u* denotes a discrete phase-state vector such as Z*θ*_*y*_(*t*), *θ*_*y*_(*t*^’^)[. Phase values were discretized into *N*_*bins*_:

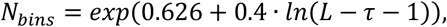

where *L* stands for number of sample points in each window, and *τ* is the lag in sample points. Finally, *dPTE* was calculated as:

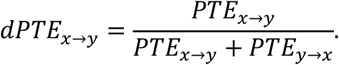

For each channel pair, a time-varying *dPTE* series was computed using a sliding window approach, with a window size of 30 s and a step size of 15 s. The median value of the resulting time series was taken as the *dPTE* of that channel pair.

The computed *dPTE* values were then used to construct the directed weighted connectivity matrix to infer the plausible sources and sinks in alpha-weaved network. For a given channel pair, the connectivity weight in each stage was defined as “median *dPTE* − 0.5”. Each channel’s specific anatomical location (e.g. inferior frontal) was mapped to a general region (e.g. frontal); thus, the weight between two general regions was defined as the averaged weights of all the belonging channel pairs. The “out-strength” for a general region was the sum of the all its output weights (i.e., positive *dPTE* − 0.5 values when the region was defined as the source). The in-strength of a given region was defined as the absolute sum of all incoming weights (i.e., negative *dPTE* − 0.5 values when the region was defined as the source). Then the net connectivity magnitude for a given region was calculated as its out-strength subtracted by its in-strength. General regions containing less than 5 channels were excluded from the analysis.

### Slow-wave activity

Signals were band-pass filtered between 0.2 and 2 Hz to isolate slow-wave activity. We defined the interval between two consecutive zero crossings as a half-wave. For each half-wave, we calculated its duration and peak amplitude (i.e., the maximum absolute amplitude). To estimate time-varying slow-wave properties, we applied a sliding-window approach using a window length of 60 s and a step size of 6 s. Within each window, (1) peak amplitude was defined as the mean peak amplitude of all half-waves contained within the window; (2) half-wave duration was defined as the mean duration of all half-waves within the window; and (3) half-wave incidence was defined as the number of half-waves occurring within the window divided by 60 s. For each subject, the median value across channels was used for all three measures.

### Phase-amplitude coupling

Signals were first band-pass filtered into the frequency bands of interest. The analytic signal was then obtained using the Hilbert transform, from which the instantaneous phase and amplitude (envelope) were extracted. For PAC analysis, we obtained the instantaneous slow-wave phase and the instantaneous alpha- or beta-band amplitude, and constructed phase-amplitude histograms using 16 phase bins spanning 0–2π. The modulation index (MI) was then computed to quantify phase-amplitude coupling using a previously established method^105^:

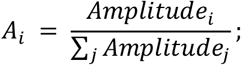

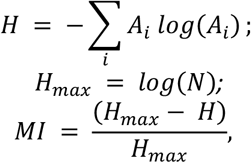

where Amplitude_*i*_ denotes the mean amplitude in the _*i*_-th phase bin, and *A*_*i*_ denotes the corresponding normalized amplitude. *H* represents the Shannon entropy for the normalized amplitude vector *A*, and *N* denotes the total number of bins (i.e., *N* = 16).

To evaluate the significance of the observed MI, we conducted a permutation-based shufle test. The observed MI was calculated by constructing a phase-amplitude histogram from the phase-amplitude pairs and computing the MI from the resulting distribution. To generate a null distribution, the amplitude time series was randomly circularly shifted relative to the phase time series across *n*_*shuffle*_ = 1000 iterations, thereby disrupting the observed PAC while preserving the internal structure of each signal. The MI was recalculated for each shufled instance, and the resulting values were used to construct the null distribution. The *p*-value was defined as the proportion of shufled MI values that were greater than or equal to the observed MI.

To investigate temporal changes in PAC, we computed the MI and corresponding phase-amplitude histograms using a sliding window with a length of 30 s and a step size of 6 s. A channel was considered to exhibit significant PAC during Anesthesia if at least 70% of the time windows within that stage yielded a *p*-value below 0.05. For these channels, the median MI within each stage was used for cross-stage comparisons using the two-tailed Wilcoxon signed-rank test.

To examine changes in the preferred phase of PAC, we calculated the complex average:

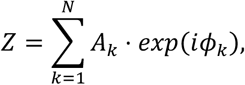

where *N* denotes the total number of phase bins, *A*_*k*_ denotes the normalized amplitude (i.e., the value of the phase-amplitude histogram) in the *k*-th phase bin, and *ϕ*_*k*_ denotes the center phase of the *k*-th bin. Here, _*i*_ is the imaginary unit. The preferred phase was then computed as:

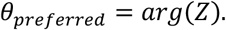

To assess how the preferred phase changes across stages for each channel, we first computed the mean phase-amplitude histogram over time during Anesthesia and TE respectively. From these averaged histograms, we extracted the preferred phase in each stage and calculated the difference between the two phases.

### Bayesian state-space model for behavioral modelling

We used a Bayesian state-space model to estimate the time-varying probability and credible interval for performance scores. For behavioral responses, the degree of completion may vary. For example, in a MC task, there were three possible conditions: (1) no observable action at all; (2) action initiated but incomplete; (3) complete fist clenching. Accordingly, we decomposed each task into different components.

MC components: (1) whether action initiated; (2) whether action completed.

VC components: (1) whether there was vocalization; (2) whether year of birth was correctly said; (3) whether month of birth was correctly said; (4) whether the date of birth was correctly said.

VR components: (1) whether there was vocalization; (2) whether the general category (e.g., animal, furniture) was correctly identified; (3) whether the specific object was correctly identified.

FO components: (1) whether action initiated; 2) whether action completed.

Each component was modeled as a binomial (Bernoulli) process that was time-varying. To align data from different subjects, we rescaled the time so that EoA = 0 and EoB = 1. For each component, binned successes were modeled using a binomial likelihood. The latent success probability was modeled as a Gaussian random walk in logit space, with an initial state z_*k*,1_~*N*(−10,1) and innovation scale *σ*_*k*_~*HalfNormal*(0.05) for component *k*. The latent state was transformed to probability space using the logistic function. Posterior inference was performed in PyMC^106^ using the No-U-Turn Sampler^107^, with 5000 posterior draws, 2000 tuning iterations, and target_accept=0.99. Then we calculated the final score at each time point. The expected score was the average of all the component probabilities:

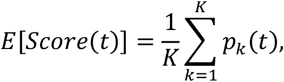

where *K* is the total number of components, and *p*_*k*_ is the probability of success for component *k* at time *t*.

### Rate of behavioral recovery

To assess individual variability in the recovery of conscious behaviors, we used the number of blocks (*t*) required to reach consistently perfect performance as an index of recovery rate for each task. For MC, VC, and FO, *t* was defined as the first block of five consecutive blocks with perfect scores. For VR, it was defined as the first block of four consecutive blocks with perfect scores, given the slower recovery of visual function. We tested whether patients differed in recovery rate by comparing the observed between-patient variance in *t* with a null distribution generated under the hypothesis that all patients shared an identical recovery rate. Specifically, under the null model, we fit a shared logistic function (block) by pooling data across all patients, and then performed 5,000 parametric bootstrap simulations while preserving each patient’s original number of blocks. For each simulated dataset, we recomputed each patient’s *t* for consistently perfect performance and calculated the between-patient variance, *Va*(*t*), thereby generating a null distribution. Monte Carlo *p*-values were computed as the proportion of bootstrap variances greater than or equal to the observed variance.

### Reaction time of verbal responses

To automatically determine participants’ speech onset times, we employed an unsupervised K-means clustering approach on their vocal responses. All audio recordings were initially low-pass filtered using a finite impulse response (FIR) filter (passband: 2600 Hz; stopband: 2800 Hz; passband ripple: 1 dB; stopband attenuation: 60 dB; sampling rate: 8192 Hz) to remove high-frequency noise. Filtered signals were z-scored to normalize amplitude across trials and subjects. Next, a sliding window analysis was conducted over the 1.5–7.0 s post-stimulus interval, with a window length of 60 ms and a step size of 10 ms. For each window, four time-domain acoustic features were computed: standard deviation, energy, maximum amplitude, and zero-crossing rate. Standard deviation reflects signal variability. Energy was calculated as the sum of squared amplitudes, reflecting the overall intensity of signals. Maximum amplitude identifies sudden intensity peaks. Zero-crossing rate indicates how quickly the waveform alternates in polarity. These features captured both amplitude- and frequency-domain properties relevant to speech detection. K-means clustering (*k* = 2) was performed independently for each trial to classify temporal windows as either background noise or speech activity. Reaction time was defined as the earliest onset of sustained speech activity following background noise.

### Task and modality selectivity

We applied a sliding-window multitaper method to compute the high-gamma band (80–150 Hz) power for each channel demonstrating task or modality selectivity, using “dpss” tapers, a 1 Hz frequency step, a 0.3 s time window, and a 10 ms step size. The power was then averaged across all frequency steps. Task and modality selectivity was determined through visual inspection of (1) averaged power during different tasks and rest; (2) raster plots showing power in each trial. Visual inspection was used here due to the complexity of the signals. For auditory selectivity, high-gamma responses ought to be higher than during rest, exhibiting either a transient increase at stimulus onset (e.g. **Figure S8:8**) or a sustained elevation throughout the auditory stimulus (e.g. **Figure S8:34**). A channel was classified as MC-selective if high-gamma power was maximal during MC trials relative to all other task conditions. Same logic applied to VR and FO selectivity. Verbal selectivity should reveal salient high-gamma increases after stimulus offset during both VC and VR trials, but not during others. Since visual selectivity (VR) demonstrated simpler patterns than others, an additional statistical criterion was applied to validate VR selectivity. High-gamma power was compared across four tasks during the 100–700 ms after stimulus onset. For each channel, we first performed one-tailed Mann–Whitney U tests (α = 0.05) comparing responses during VR trials with those during each of the other task conditions (MC, VC, and FO). Subsequently, we computed Cliff’s delta effect sizes for all VR-versus-other comparisons to quantify practical significance, requiring an effect size greater than 0.28 to indicate a meaningful difference. All VR-selective channels identified through visual inspection met both criteria.

### High-gamma response strength

We applied a sliding-window multitaper method to compute the high-gamma band (80–150 Hz) power for each channel demonstrating task or modality selectivity, using 7 tapers, a 2 Hz frequency step, a 0.3 s time window, and a 2 ms step size. The power was then averaged across all frequency steps. Given that both the behavioral and neural responses to FO were relatively longer and less aligned to stimulus onset or offset, the response strength for FO-selective channels were determined differentially. First, the critical window was identified for each type of selectivity using a sliding search approach. For all selectivity except for FO, we calculated the mean power across all trials under each selectivity during FW, and the time window exhibiting the highest mean power was identified as the critical window. The duration of the time window for each selectivity, listed in **Table 1**, was decided based on the characteristics of corresponding high-gamma responses. For FO, the critical window was determined trial by trial, independent of the data during FW. The start and end of the search are specified in **Table 1**.

**Table 1.** Parameters for calculating response strength.

| Selectivity | CW duration | Search range |
| --- | --- | --- |
| Audition (all tasks) | 0.25 s | Instruction onset – offset |
| Verbalization (VC+VR) | 1 s | 1 s before instruction offset – trial end |
| MC | 2 s | 1 s before instruction offset – trial end |
| VR | 0.25 s | Instruction onset – offset |
| FO | 4 s | 1 s before instruction offset – trial end |

For each trial, the response strength was calculated as follows:

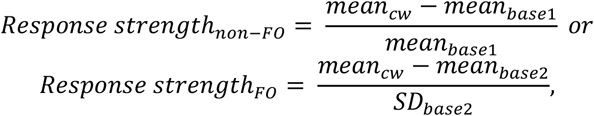

where *mean*_*c*w_ stands for the mean power during the critical window; *mean*_*base*1_ is the mean power during 400 ms before instruction onset; *mean*_*base*2_ is the mean power during 5 seconds before instruction offset; *SD*_*base*2_ is the standard deviation during 5 seconds before instruction offset. Finally, for each type of selectivity under each stage, a channel’s response strength was calculated as the median response strength across all trials. One-tailed Wilcoxon signed-rank test between stages was performed to determine where there was a significant progressive increase across stages. Two-tailed Wilcoxon rank-sum test was used to test if there was a significant difference in response strength between channels showing selectivity versus no selectivity during Task Engagement.

## Acknowledgement

We thank Glenn Tam, Luming Yang, Jiemin Jia, Haiyan Wu, Junheng Li, Yi Sun, Ren Sun, Tian Xu, Chong Chen, Bo Li, and Xiaoming Zhang for insightful discussions and feedback on the project; Xinjing Li and Hao Zhu for guiding equipment setup; Xing Tian for sharing equipment; and all patients and their families for their generosity to support this study. The research was funded by the Westlake Fellows Program at Westlake University and Funding No. JCYJ20240813141105007 supported by Shenzhen Science and Technology Innovation Committee, China.

## Declaration of generative AI and AI-assisted technologies

During the preparation of this work, the authors used ChatGPT and DeepSeek for programming assistance and language editing. No scientific ideas, interpretations, analyses, results, or conclusions were generated by artificial intelligence. The authors reviewed and edited the output as needed and take full responsibility for the content of the published article.

