## Supplementary information for "A human intracranial map of consciousness returning from anesthesia"

**Table S1**

| General region | Specific region | Channel count |
|---|---|---|
| basal ganglia | putamen | 19 |
| | Pallidum | 2 |
| cingulate | caudalanteriorcingulate | 4 |
| | isthmuscingulate | 4 |
| | rostralanteriorcingulate | 4 |
| | posteriorcingulate | 1 |
| diencephalon | Thalamus-Proper | 5 |
| | VentralDC | 5 |
| frontal | superiorfrontal | 47 |
| | rostralmiddlefrontal | 27 |
| | medialorbitofrontal | 26 |
| | parstriangularis | 25 |
| | lateralorbitofrontal | 18 |
| | precentral | 15 |
| | parsorbitalis | 11 |
| | paracentral | 7 |
| | parsopercularis | 5 |
| | frontalpole | 4 |
| | caudalmiddlefrontal | 2 |
| insula | insula | 76 |
| occipital | lateraloccipital | 4 |
| | pericalcarine | 3 |
| | cuneus | 2 |
| parietal | supramarginal | 25 |
| | inferiorparietal | 18 |
| | superiorparietal | 14 |
| | precuneus | 11 |
| | postcentral | 9 |
| temporal | hippocampus | 57 |
| | amygdala | 33 |
| | middletemporal | 30 |
| | superiortemporal | 19 |
| | parahippocampal | 8 |
| | fusiform | 5 |
| | bankssts | 4 |
| | inferiortemporal | 4 |
| | transversetemporal | 3 |
| | **Grand total** | **556** |

**Table S1. Anatomical locations of all channels recorded in the gray matter.**

**Table S2**

| Nearest general region | Nearest specific region | Channel count |
|---|---|---|
| basal ganglia | putamen | 15 |
| | accumbens | 1 |
| cingulate | caudalanteriorcingulate | 10 |
| | rostralanteriorcingulate | 7 |
| | isthmuscingulate | 3 |
| | posteriorcingulate | 3 |
| diencephalon | thalamus | 5 |
| frontal | superiorfrontal | 62 |
| | parstriangularis | 43 |
| | lateralorbitofrontal | 41 |
| | rostralmiddlefrontal | 38 |
| | medialorbitofrontal | 18 |
| | precentral | 15 |
| | parsorbitalis | 7 |
| | caudalmiddlefrontal | 5 |
| | paracentral | 5 |
| | parsopercularis | 4 |
| | frontalpole | 3 |
| insula | insula | 50 |
| occipital | pericalcarine | 10 |
| | lateraloccipital | 5 |
| | cuneus | 2 |
| | lingual | 1 |
| parietal | supramarginal | 27 |
| | inferiorparietal | 18 |
| | precuneus | 18 |
| | postcentral | 14 |
| | superiorparietal | 10 |
| temporal | middletemporal | 77 |
| | superiortemporal | 41 |
| | inferiortemporal | 27 |
| | fusiform | 13 |
| | hippocampus | 12 |
| | parahippocampal | 10 |
| | amygdala | 9 |
| | transversetemporal | 5 |
| **sum** | | **634** |

**Table S2. Anatomical locations of all channels recorded in the white matter.** Locations indicate the nearest gray matter regions.

**Table S3**

| Sub | Age | Gender | N block total | N stage 1 | N stage 2 | N stage 3 | N pre-extubation | N post-extubation | EoA ~ EoB (min) |
|---|---|---|---|---|---|---|---|---|---|
| 1 | 27 | f | 10 | 0 | 3 | 7 | 3 | 7 | 30.55 |
| 2 | 17 | f | 32 | 0 | 19 | 13 | 18 | 14 | 36.1 |
| 3 | 42 | f | 27 | 10 | 14 | 3 | 23 | 4 | 18.43 |
| 4 | 46 | f | 29 | 10 | 7 | 12 | 16 | 13 | 9.46 |
| 5 | 33 | m | 39 | 10 | 12 | 17 | 18 | 21 | 17.31 |
| 6 | 32 | m | 47 | 10 | 23 | 14 | 33 | 14 | 25.68 |
| 7 | 19 | f | 34 | 3 | 19 | 12 | 20 | 14 | 21.6 |
| 8 | 28 | m | 45 | 10 | 15 | 20 | 25 | 20 | 15.38 |
| 9 | 35 | f | 37 | 10 | 11 | 16 | 21 | 16 | 12.55 |
| 10 | 21 | m | 30 | 10 | 9 | 11 | 19 | 11 | 11.25 |

**Table S3: Subject general information.** Patients completed unequal total numbers of blocks due to differential recovery paces. Stage 1 = Anesthesia; Stage 2 = Transition; Stage 3 = Task Engagement. Ideally, Stage 1 lasted for 10 blocks and Stage 3 included 10 continuous blocks of perfect performance. "N block total" = "N stage 1" + "N stage 2" + "N stage 3" = "N pre-Extubation" + "N post-Extubation". The last column lists the duration from the end of anesthesia to the emergence of behaviors in each subject.

### Figure S1

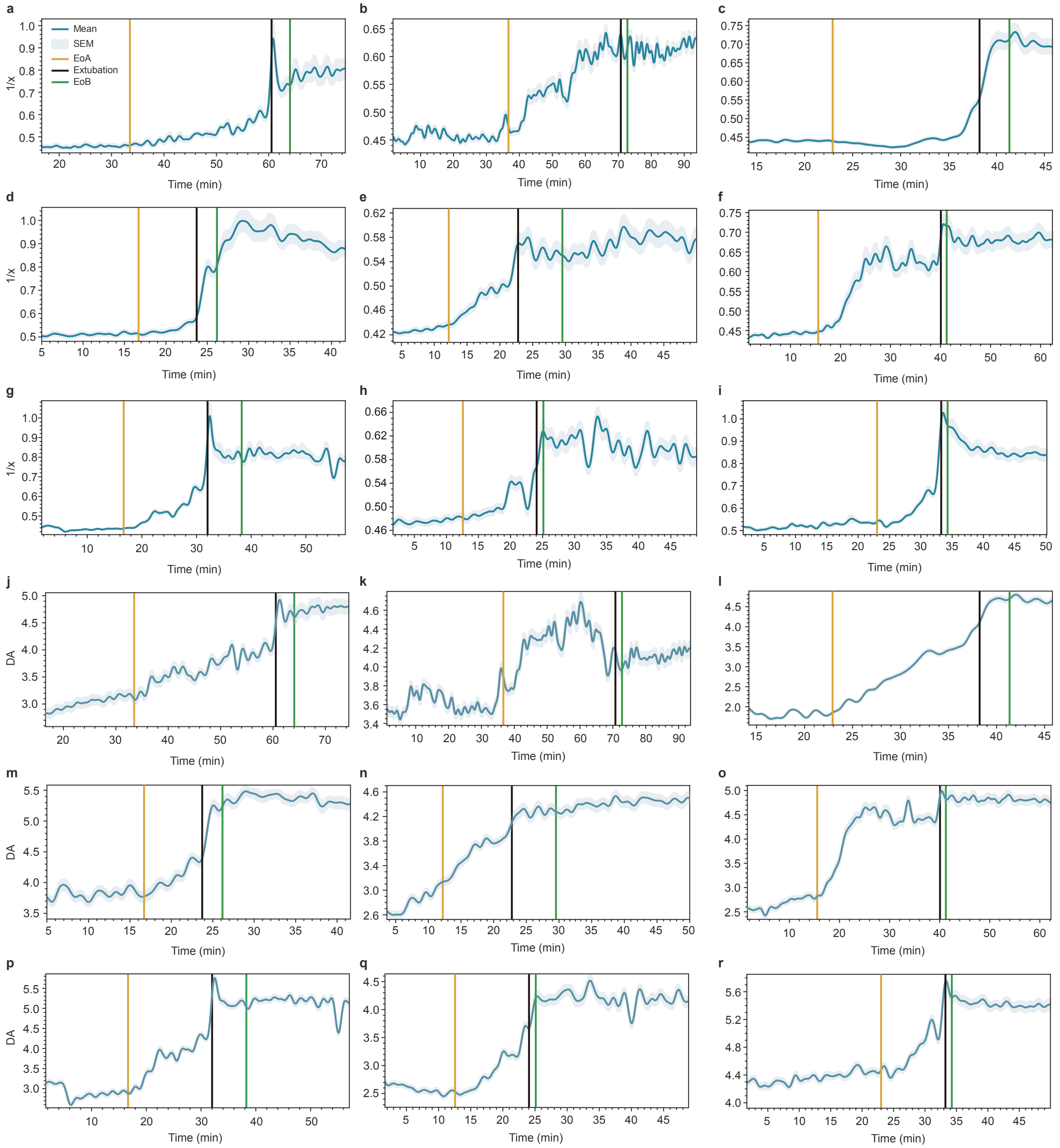


**Figure S1 | E/I and $DA$ change in each individual.** Each panel shows the E/I (**a–i**) or $DA$ (**j–r**) progression in one subject. Main line represents the mean $1/x$ of all the channels in that subject. Shaded regions signify SEM. EoA = End of Anesthesia; EoB = Emergence of Behaviors.

### Figure S2

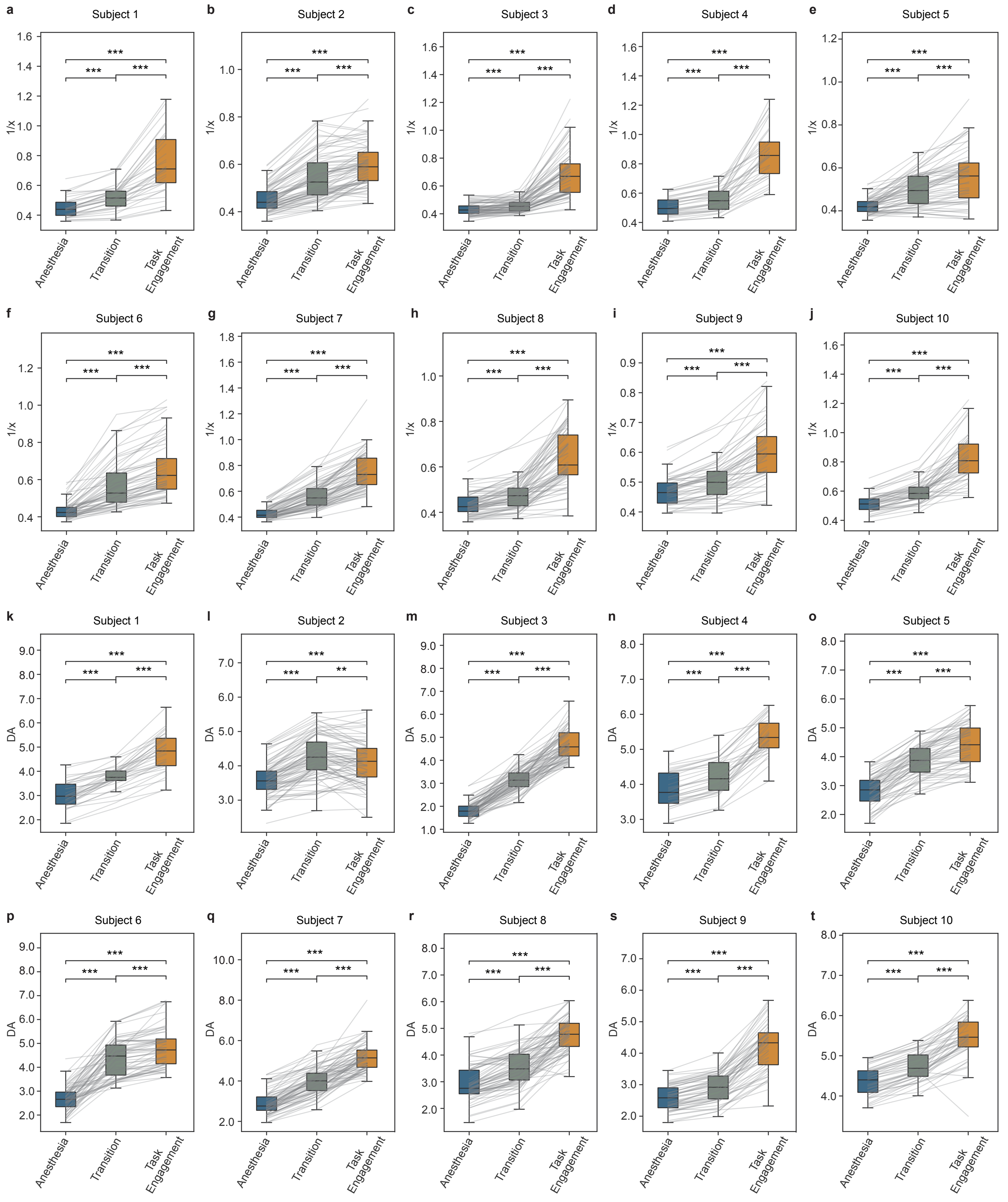


**Figure S2 | E/I and $DA$ continuously increased across stages during the recovery from anesthesia.** Each panel shows how E/I (**a–j**) or $DA$ (**k–t**) evolved in each individual. Each line represents data from one channel. Pairwise comparison was performed with signed-rank test.

### Figure S3

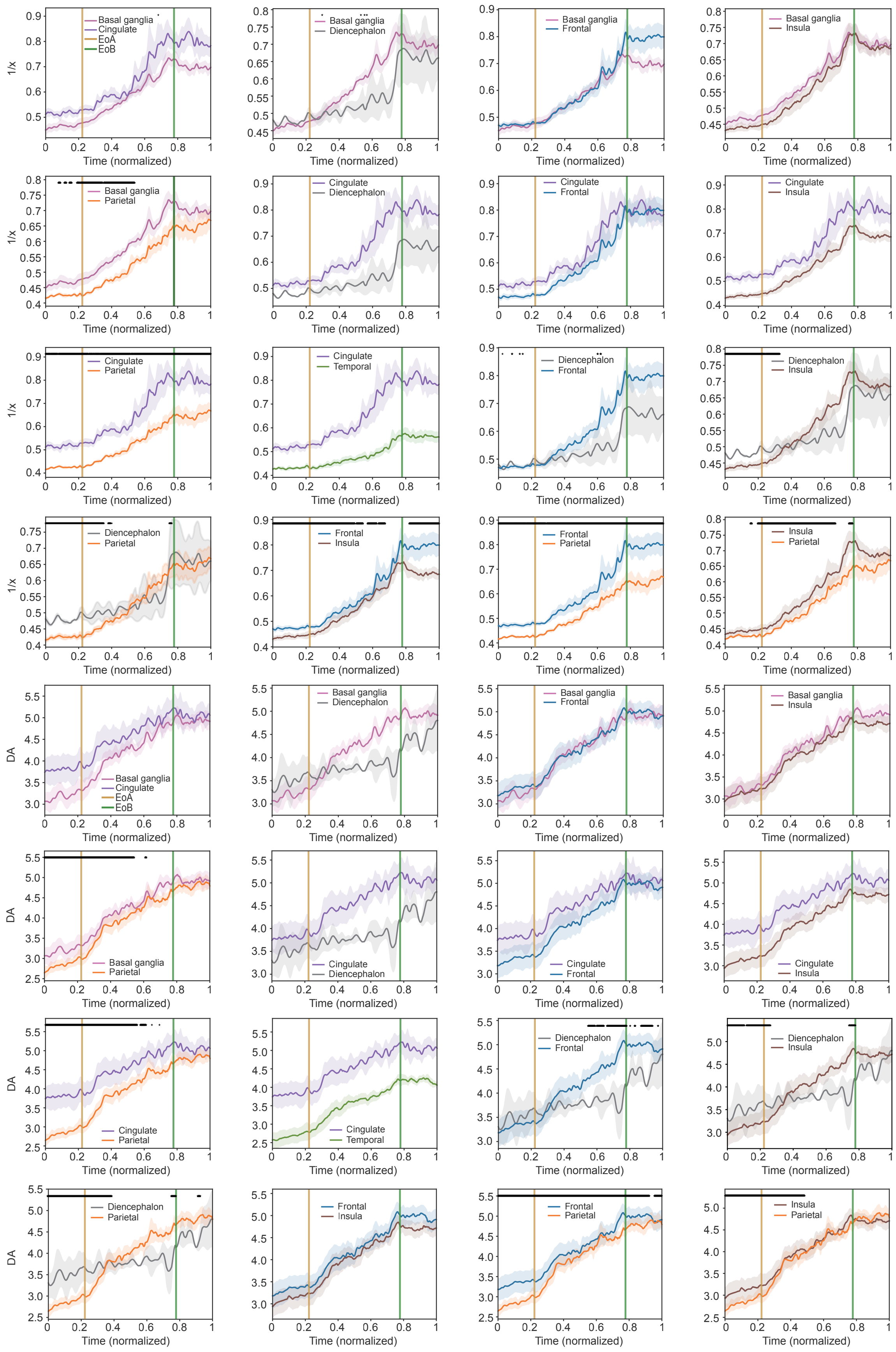
Basal ganglia
Cingulate
EoA
EoB
Diencephalon
Frontal
Insula
Parietal
Temporal
1/x
DA
Time (normalized)

### Figure S3 caption

**Figure S3 | Comparison of E/I and $DA$ across brain regions.** Each panel shows pairwise comparison of $1/x$ or $DA$ between two general regions. Time courses were temporally aligned to juxtapose data from multiple subjects on a same time axis. Black horizontal lines indicate the time points that showed significant difference between regions (linear mixed model, FDR-corrected $p$ < 0.05). Shaded regions represent SEM.

### Figure S4

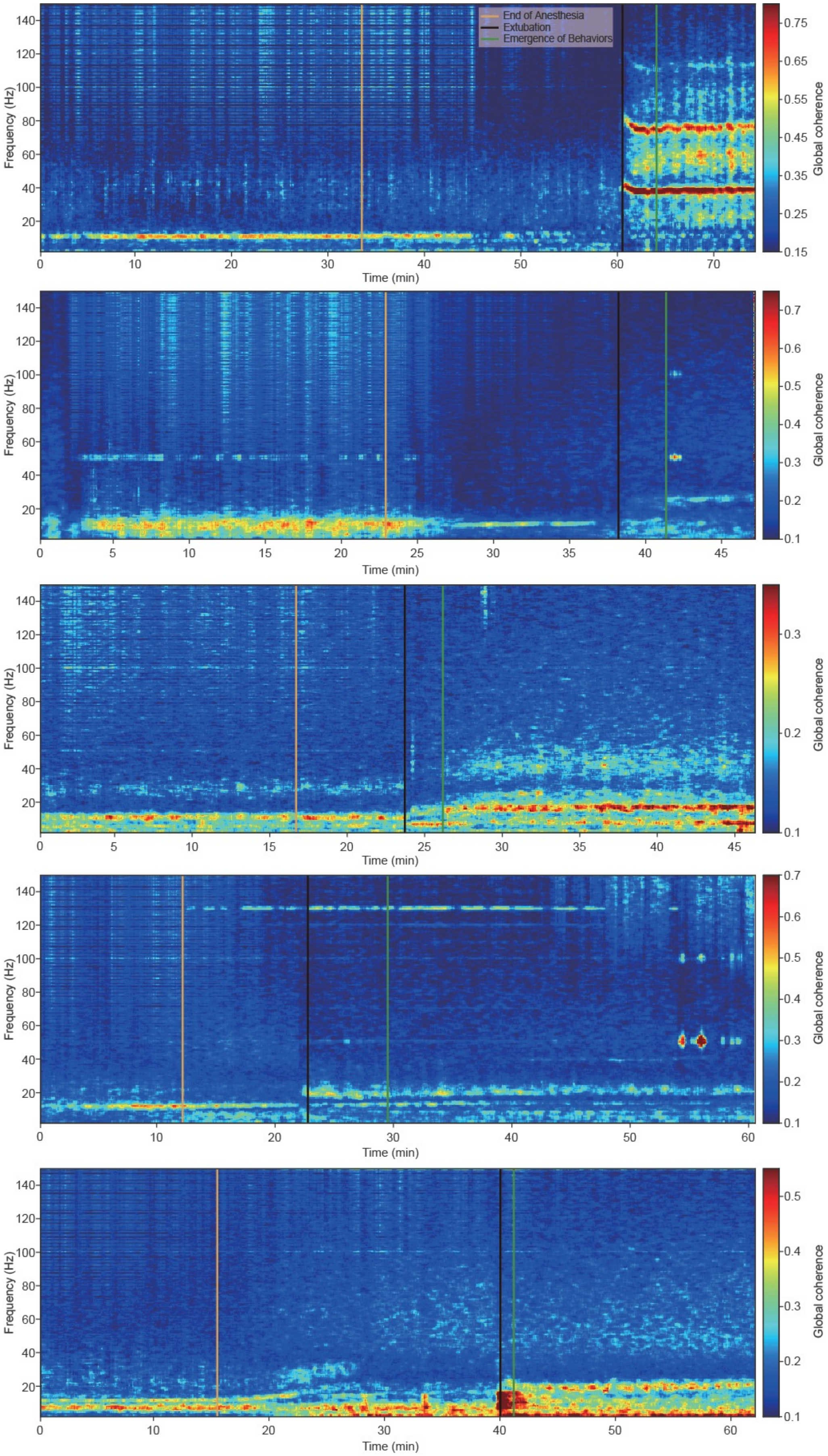
End of Anesthesia
Extubation
Emergence of Behaviors
Frequency (Hz)
Time (min)
Global coherence

### Figure S4 continued

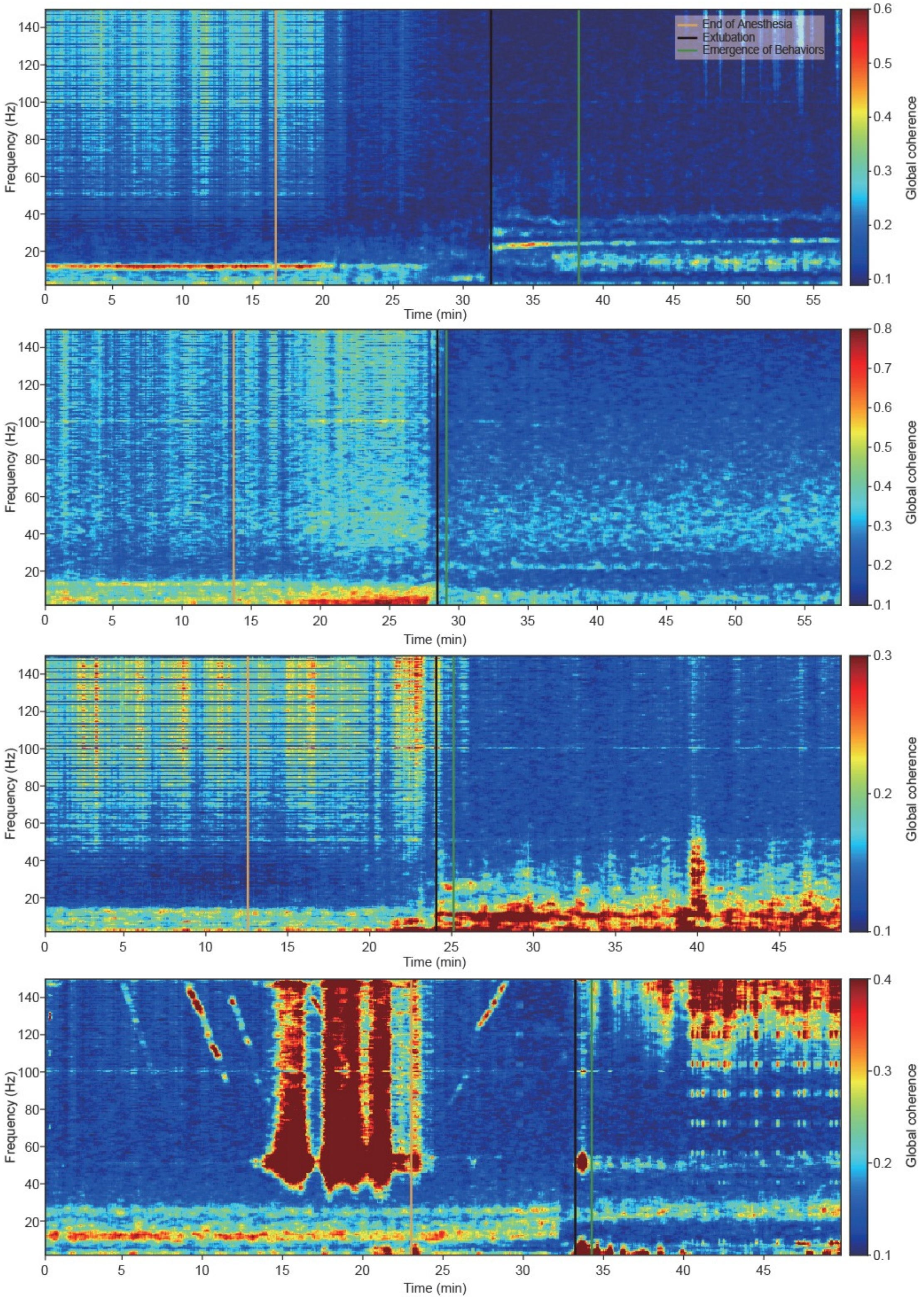


**Figure S4 | Profile of global coherence in each individual.** Each panel shows data from one subject.

### Figure S5

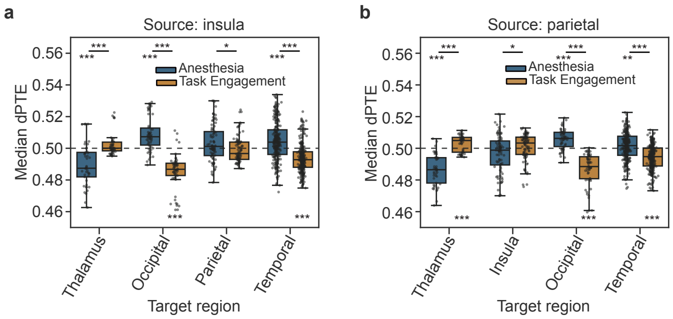


**Figure S5 | Alpha $dPTE$ from insula and parietal lobe. a.** Comparison of $dPTE$ between Anesthesia and Task Engagement, where the source was defined as the insula. **b.** Comparison of $dPTE$ between Anesthesia and TE, where the source was defined as the parietal lobe. Signed-rank test was implemented for comparison between stages and also comparison with the baseline. Regions less than 5 channels were not included in this analysis. Figure format follows that in **Figure 5m**.

### Figure S6

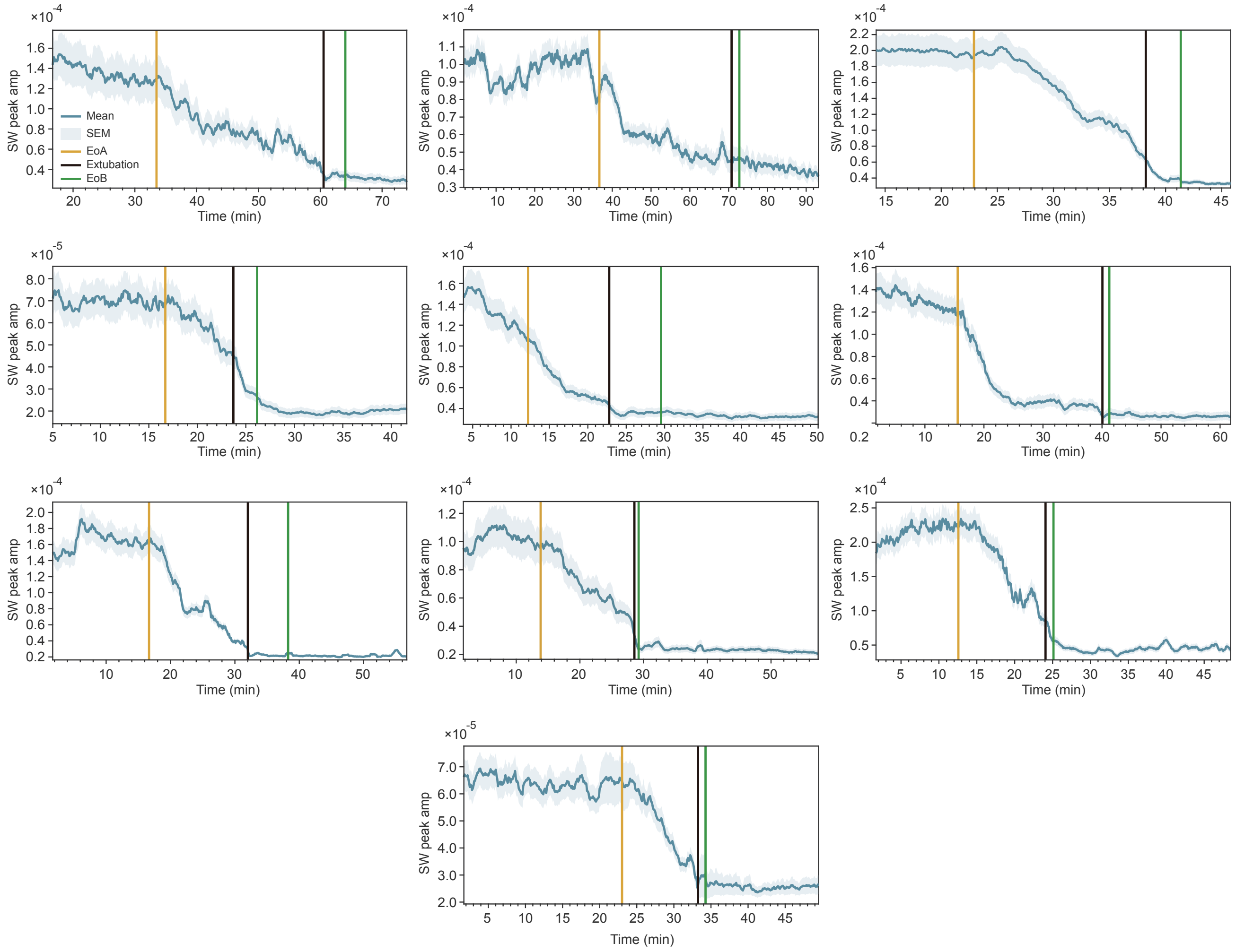


**Figure S6 | Change in slow-wave peak amplitude of each individual.** Each panel shows the SW (0.2–2 Hz) peak amplitudes in one subject. Main line represents the mean amplitude of all the channels in that subject. Shaded regions signify SEM. EoA = End of Anesthesia; EoB = Emergence of Behaviors.

### Figure S7

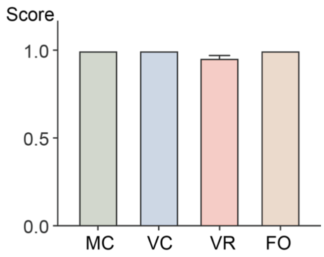


**Figure S7 | Perfect task performance during Full Wakefulness.** A score of 0 indicates no observation of any behavior, and 1 represents perfect completion. 0.5 indicates wrong or incomplete behavioral output. MC = Movement to Command, VC = Verbal Communication, VR = Visual Recognition, FO = Functional Object Use. Bar heights indicate the average scores across all subjects. The error bar signifies SEM.

### Figure S8

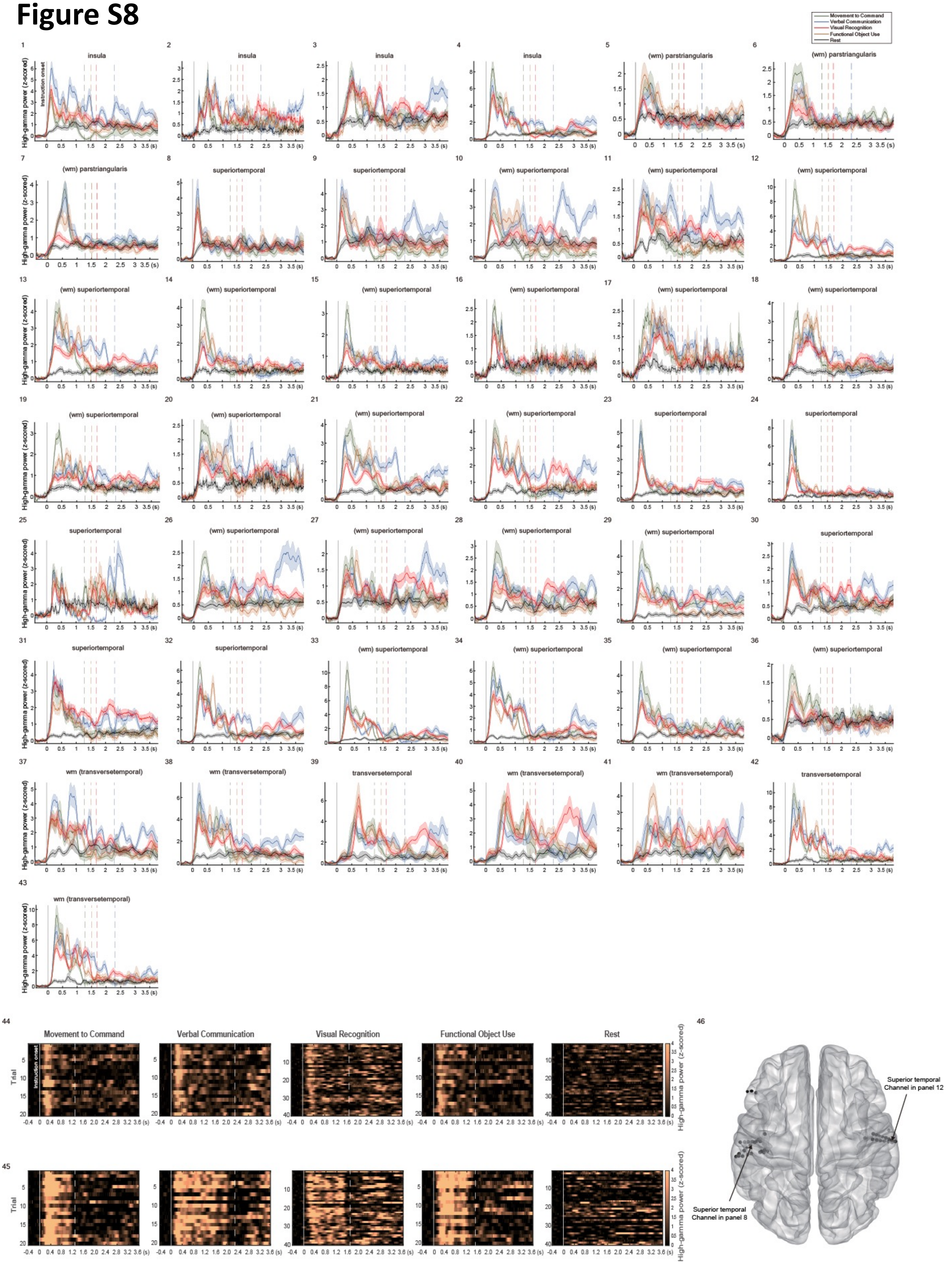

Movement to Command
Verbal Communication
Visual Recognition
Functional Object Use
Rest
insula
(wm) parstriangularis
superiortemporal
(wm) superiortemporal
wm (transversetemporal)
transversetemporal
High-gamma power (z-scored)
Instruction onset
Trial
Superior temporal Channel in panel 12
Superior temporal Channel in panel 8

### Figure S8 caption

**Figure S8 | All auditory-selective channels.**
**1–43.** Each panel illustrates the z-scored high-gamma power in response to each task during FW for one auditory-selective channel. Shaded regions indicate SEM.
**44**. Raster plot showing the high-gamma power in all trials from the example channel in panel **8**.
**45**. Raster plot showing the high-gamma power in all trials from the example channel in panel **12**. Responses in **44** are transient, while those in **45** are sustained.
**46**. Locations of all auditory-selective channels. Arrows point to the two example channels in **44–45**.

### Figure S9

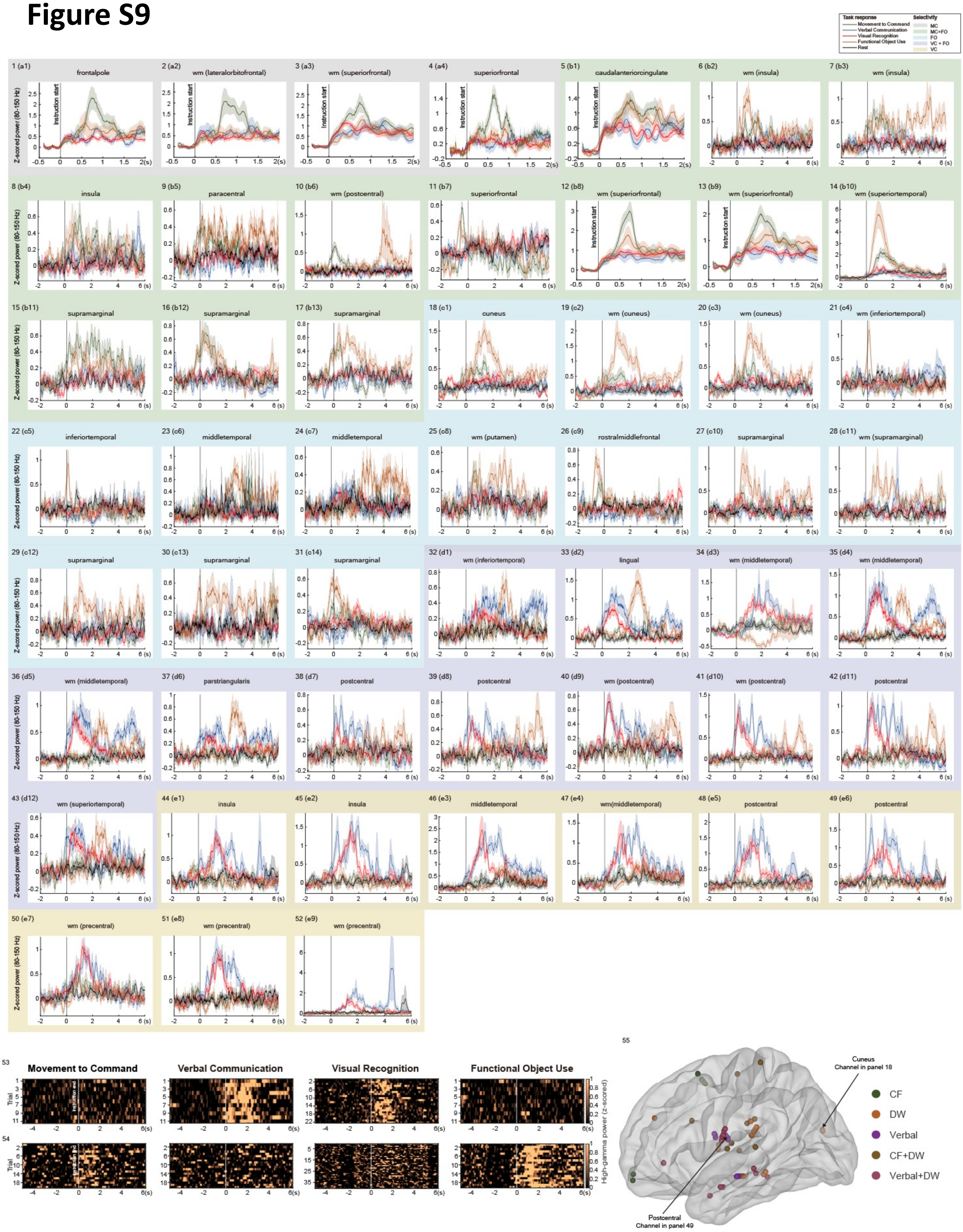

Task response
Movement to Command
Verbal Communication
Visual Recognition
Functional Object Use
Rest
Selectivity
MC
MC+FO
FO
VC + FO
VC
1 (a1) frontalpole
2 (a2) wm (lateralorbitofrontal)
3 (a3) wm (superiorfrontal)
4 (a4) superiorfrontal
5 (b1) caudalanteriorcingulate
6 (b2) wm (insula)
7 (b3) wm (insula)
8 (b4) insula
9 (b5) paracentral
10 (b6) wm (postcentral)
11 (b7) superiorfrontal
12 (b8) wm (superiorfrontal)
13 (b9) wm (superiorfrontal)
14 (b10) wm (superiortemporal)
15 (b11) supramarginal
16 (b12) supramarginal
17 (b13) supramarginal
18 (c1) cuneus
19 (c2) wm (cuneus)
20 (c3) wm (cuneus)
21 (c4) wm (inferiortemporal)
22 (c5) inferiortemporal
23 (c6) middletemporal
24 (c7) middletemporal
25 (c8) wm (putamen)
26 (c9) rostralmiddlefrontal
27 (c10) supramarginal
28 (c11) wm (supramarginal)
29 (c12) supramarginal
30 (c13) supramarginal
31 (c14) supramarginal
32 (d1) wm (inferiortemporal)
33 (d2) lingual
34 (d3) wm (middletemporal)
35 (d4) wm (middletemporal)
36 (d5) wm (middletemporal)
37 (d6) parstriangularis
38 (d7) postcentral
39 (d8) postcentral
40 (d9) wm (postcentral)
41 (d10) wm (postcentral)
42 (d11) postcentral
43 (d12) wm (superiortemporal)
44 (e1) insula
45 (e2) insula
46 (e3) middletemporal
47 (e4) wm(middletemporal)
48 (e5) postcentral
49 (e6) postcentral
50 (e7) wm (precentral)
51 (e8) wm (precentral)
52 (e9) wm (precentral)
Z-scored power (80-150 Hz)
Instruction start
53
54
Movement to Command
Verbal Communication
Visual Recognition
Functional Object Use
Trial
High-gamma power (z-scored)
55
Cuneus
Channel in panel 18
Postcentral
Channel in panel 49
CF
DW
Verbal
CF+DW
Verbal+DW

### Figure S9 caption

**Figure S9 | All motor- and verbal-selective channels.**
**1–52.** Each panel illustrates the z-scored high-gamma power in response to each task during Full Wakefulness for one channel. Shaded regions indicate SEM. Unless labeled “instruction start” in the panel, the power time course was aligned to instruction offset.
**53**. Raster plot showing the high-gamma power in all trials from the example channel in panel **18**.
**54**. Raster plot showing the high-gamma power in all trials from the example channel in panel **49**.
**55**. Locations of all motor- and verbal-selective channels. Arrows point to the two example channels in **53–54**.

### Figure S10

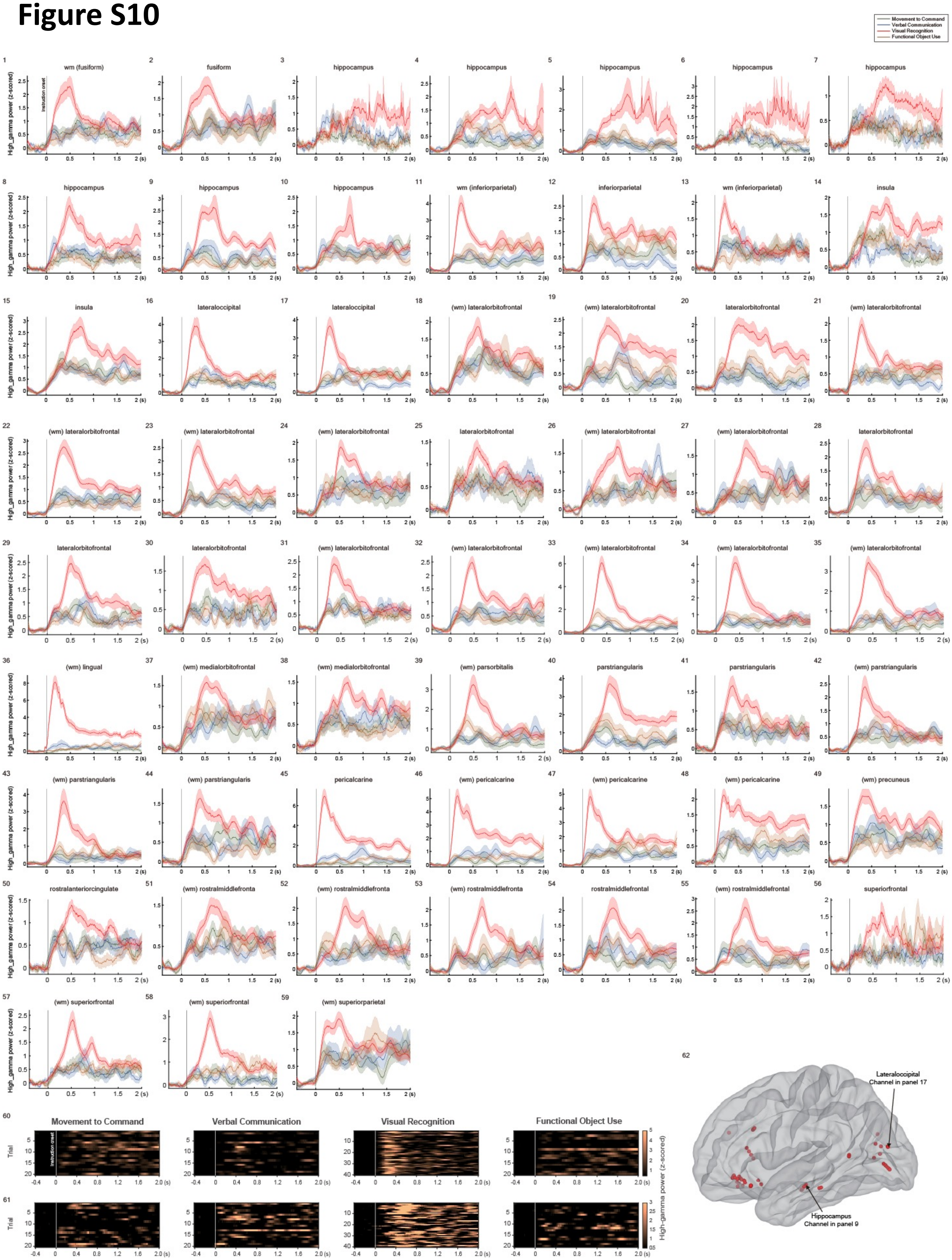

Movement to Command
Verbal Communication
Visual Recognition
Functional Object Use
High_gamma power (z-scored)
Instruction onset
1 wm (fusiform)
2 fusiform
3 hippocampus
4 hippocampus
5 hippocampus
6 hippocampus
7 hippocampus
8 hippocampus
9 hippocampus
10 hippocampus
11 wm (inferiorparietal)
12 inferiorparietal
13 wm (inferiorparietal)
14 insula
15 insula
16 lateraloccipital
17 lateraloccipital
18 (wm) lateralorbitofrontal
19 (wm) lateralorbitofrontal
20 lateralorbitofrontal
21 (wm) lateralorbitofrontal
22 (wm) lateralorbitofrontal
23 (wm) lateralorbitofrontal
24 (wm) lateralorbitofrontal
25 lateralorbitofrontal
26 (wm) lateralorbitofrontal
27 (wm) lateralorbitofrontal
28 lateralorbitofrontal
29 lateralorbitofrontal
30 lateralorbitofrontal
31 (wm) lateralorbitofrontal
32 (wm) lateralorbitofrontal
33 (wm) lateralorbitofrontal
34 (wm) lateralorbitofrontal
35 (wm) lateralorbitofrontal
36 (wm) lingual
37 (wm) medialorbitofrontal
38 (wm) medialorbitofrontal
39 (wm) parsorbitalis
40 parstriangularis
41 parstriangularis
42 (wm) parstriangularis
43 (wm) parstriangularis
44 (wm) parstriangularis
45 pericalcarine
46 (wm) pericalcarine
47 (wm) pericalcarine
48 (wm) pericalcarine
49 (wm) precuneus
50 rostralanteriorcingulate
51 (wm) rostralmiddlefronta
52 (wm) rostralmiddlefronta
53 (wm) rostralmiddlefronta
54 rostralmiddlefrontal
55 (wm) rostralmiddlefrontal
56 superiorfrontal
57 (wm) superiorfrontal
58 (wm) superiorfrontal
59 (wm) superiorparietal
60
61
Trial
High-gamma power (z-scored)
62
Lateraloccipital Channel in panel 17
Hippocampus Channel in panel 9

### Figure S10 caption

**Figure S10 | All VR-selective channels.**
**1–59.** Each panel illustrates the z-scored high-gamma power in response to each task during Full Wakefulness for one VR-selective channel. Shaded regions indicate SEM.
**60**. Raster plot showing the high-gamma power in all trials from the example channel in panel **17**.
**61**. Raster plot showing the high-gamma power in all trials from the example channel in panel **9**. Responses in **17** are more transient, while those in **9** are more sustained.
**62**. Locations of all VR-selective channels. Arrows point to the two example channels in **60–61**.

### Figure S11

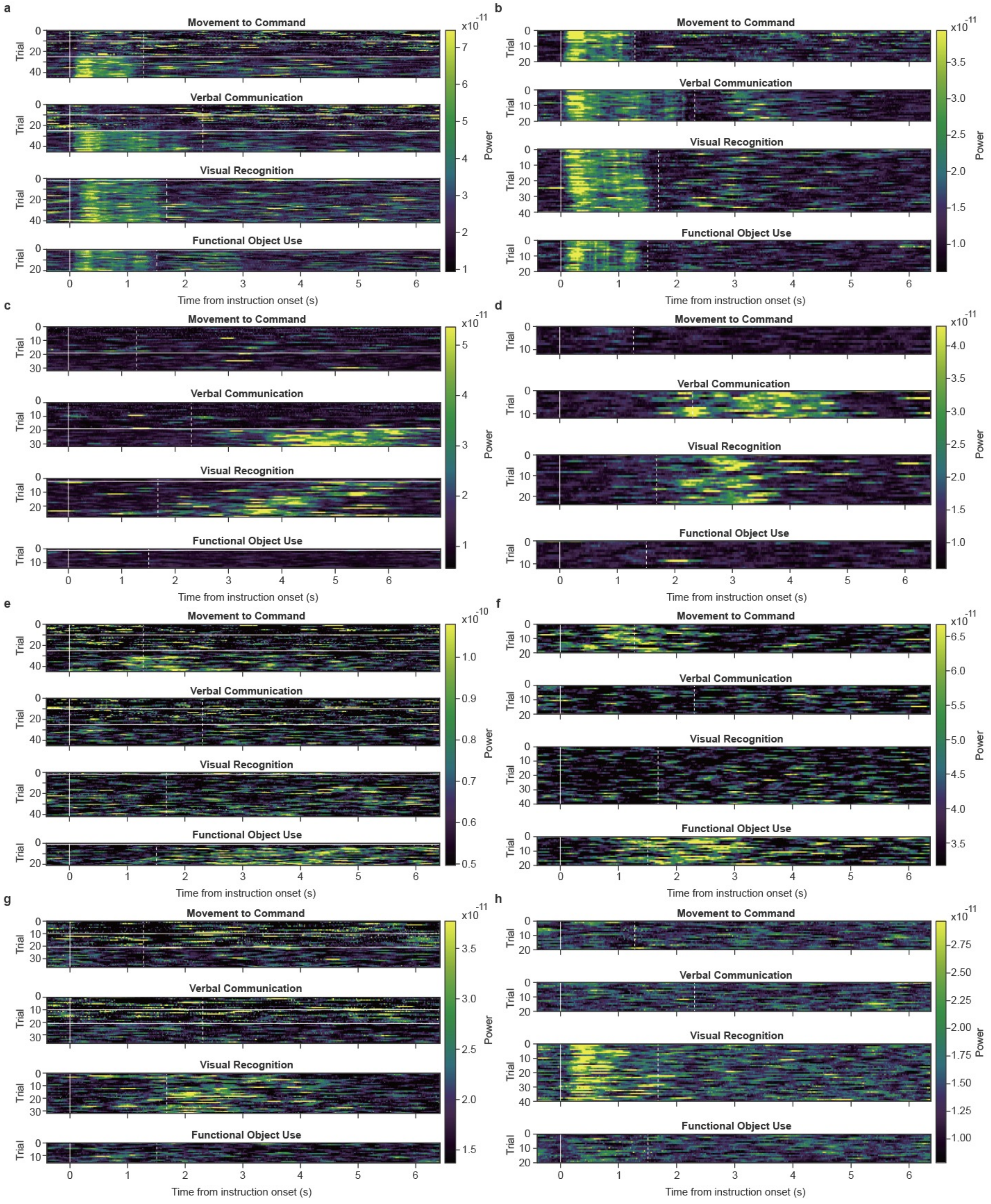
a
b
c
d
e
f
g
h
Movement to Command
Verbal Communication
Visual Recognition
Functional Object Use
Trial
Power
Time from instruction onset (s)

### Figure S11 caption

**Figure S11 | High-gamma responses became increasingly similar to those observed during FW.** Raster plots displaying the high-gamma power in all trials in the OR (left panels) and EMU (right panels), from example channels exhibiting auditory (**a–b**), verbal (**c–d**), MC+FO (**e–f**), and VR (**g–h**) selectivity. In each panel, the vertical solid line indicates instruction onset; the dashed line marks instruction offset. For responses in the OR, there could be two (EoA then EoB), one (EoB), or zero horizontal white lines, depending on the number of blocks completed in each stage and whether the task was administered only after eye opening (VR, FO).
